# Perinatal brain group 3 innate lymphoid cells are involved in the formation of murine dural lymphatics

**DOI:** 10.1101/2024.06.12.597323

**Authors:** Alba del Rio Serrato, Borja Latorre Hernández, Evie Bartl, Christina Stehle, Oliver Hölsken, Andreas Diefenbach, Chiara Romagnani, Carmen Infante-Duarte

## Abstract

The central nervous system (CNS) contains a pool of innate lymphoid cells (ILCs) of unclear composition and functionality and unknown origin. Here, we demonstrate that group 1 ILCs (inc. ex-ILC3s) and ILC2s are resident cells with low proliferative capacities and subtype specific CNS compartmentalization. We show for the first time that CNS ILC seeding and niche establishment occurs during early life and is initiated by both, ILC progenitors-like PLFZ^+^PD- 1^+^ cells and lineage committed ILCs. While group 1 ILCs and RORγt^+^ ILC3s were found within the embryonic and postnatal brain, ILC2s reached the CNS after birth and were predominantly localized within the dura mater, proving early regional distribution. Interestingly, RORγt^+^ ILC3s were only detected perinatally and vanished from the CNS as an outcome of decreased turnover and *in situ* ILC3-to-ILC1 conversion. Remarkably, we showed that perinatal RORγt^+^ ILC3s are required for the correct development of the lymphatic vessels within the dura.

## Introduction

Innate lymphoid cells (ILCs) form a heterogeneous group of tissue resident innate immune cells that act not only as key modulators of the immune response but are also involved in other processes such as tissue homeostasis, repair, and remodeling^1–4^. ILCs are categorized in five subsets based on their developmental pathways and phenotypical profiles. Group 1 ILCs includes conventional natural killer (NK) cells, that express the transcription factor (TF) eomesodermin (Eomes)^5^, and helper ILC1s. Both subsets express T-bet^5^ and produce large amounts of IFN-γ^6,7^. Group 2, characterized by high expression of GATA-3, produce amphiregulin, IL-4, IL-5, or IL-13 and are enriched in barrier structures such as the skin, lung, and intestine. Finally, RORγt-dependent group 3 ILCs, including NKp46^+/-^ ILC3s and CCR6^+^ lymphoid tissue inducer (LTi) cells, are characterized by the production of lymphotoxin-α, IL- 17 and IL-22.

ILCs develop from the common lymphoid progenitor (CLP) and appear to seed tissues already at prenatal stages, displaying organ-specific distribution and composition^8,9^. The main hematopoietic ILC sources are the fetal liver and the adult bone marrow; however, circulating fetal ILC progenitors that seed peripheral tissues and undergo *in situ* differentiation have also been observed^10–12^. Indeed, parabiotic experiments confirmed that the ILC pool in many tissues is comprised by both embryonic-derived long-lived cells and a fraction that is generated during adulthood^13–16^. Additionally, plasticity events mediated by e.g. tissue-derived cytokines have been reported within all ILC subsets^17,18^, leading to the generation of a plethora of intermediate populations^19–22^. Overall, these processes of layered ontogeny, adult-derived turnover, and ILC plasticity, facilitate the adaptation of tissue-specific ILCs to external stimuli during immune activation and physiological processes of tissue development, homeostasis and aging^12,23,24^.

ILCs have been mainly described in barrier structures such as the airways, digestive tract, and skin; though, we and others have reported on resident ILCs within the central nervous system (CNS)^6,25–27^. Most of these studies focused on the ILC distribution and function in inflamed CNS compartments. Within the meningeal barriers, ILCs modulate T cell activation and trafficking during autoimmune neuroinflammation by acting as antigen-presenting cells^28,29^, tissue healing after spinal cord injury^30^, and sex-dimorphism in the T cell response during experimental autoimmune encephalomyelitis (EAE)^31^. However, because of the low ILC numbers in the steady-state CNS, little is known on dynamics of tissue seeding, local differentiation, and their possible roles in CNS homeostasis and development.

Here, we demonstrated that group 1 and ILC2s are quiescent CNS resident cells with low proliferative capacities and compartment-specific distribution. Importantly, we defined the formation of the CNS-ILC niche during embryonic development and postnatal stages. While group 1 ILCs, RORγt^+^ ILC3s, and LTi cells were detected in the brain prenatally, ILC2s mainly reached the brain after birth. Further, the ILC distribution along the different CNS compartments is already established after the second week of life. RORγt^+^ ILC3s, transiently appeared during perinatal stages of brain development but are absent in the adult CNS due to, in part, postnatal *in situ* ILC3 to ILC1 conversion. Ex-ILC3s appeared as a population with long-lived capacities as a proportion of these cells remained in the brain until, at least, 7 months of age. Remarkably, we showed that the presence of perinatal RORγt^+^ ILC3s is required for the correct development of the lymphatic vessels within the dura mater.

## Results

### Group 1 ILCs (including ex-ILC3s) and ILC2s but not ILC3s are tissue-resident immune cells in the young adult CNS

To determine the ILC profile of the murine adult CNS, we performed high-dimensional flow cytometry and identified group 1 ILCs and ILC2s as main ILC populations within the tissue (Representative gating strategy is shown in Fig. 1A; Sup. Fig. 1A). To confirm these results, we examined an available scRNA-seq dataset of the whole murine brain and brain barrier- sorted immune cells^32^. Group 1 and 2 ILCs were identified as the sole ILC populations along the different CNS compartments (Fig. 1B), after excluding clusters with markers for other immune cell populations and selecting those clusters with the highest ILC signature score (Sup. Fig. 1B-D; ILC signature genes depicted in Sup. Table 1).

**Figure 1.**
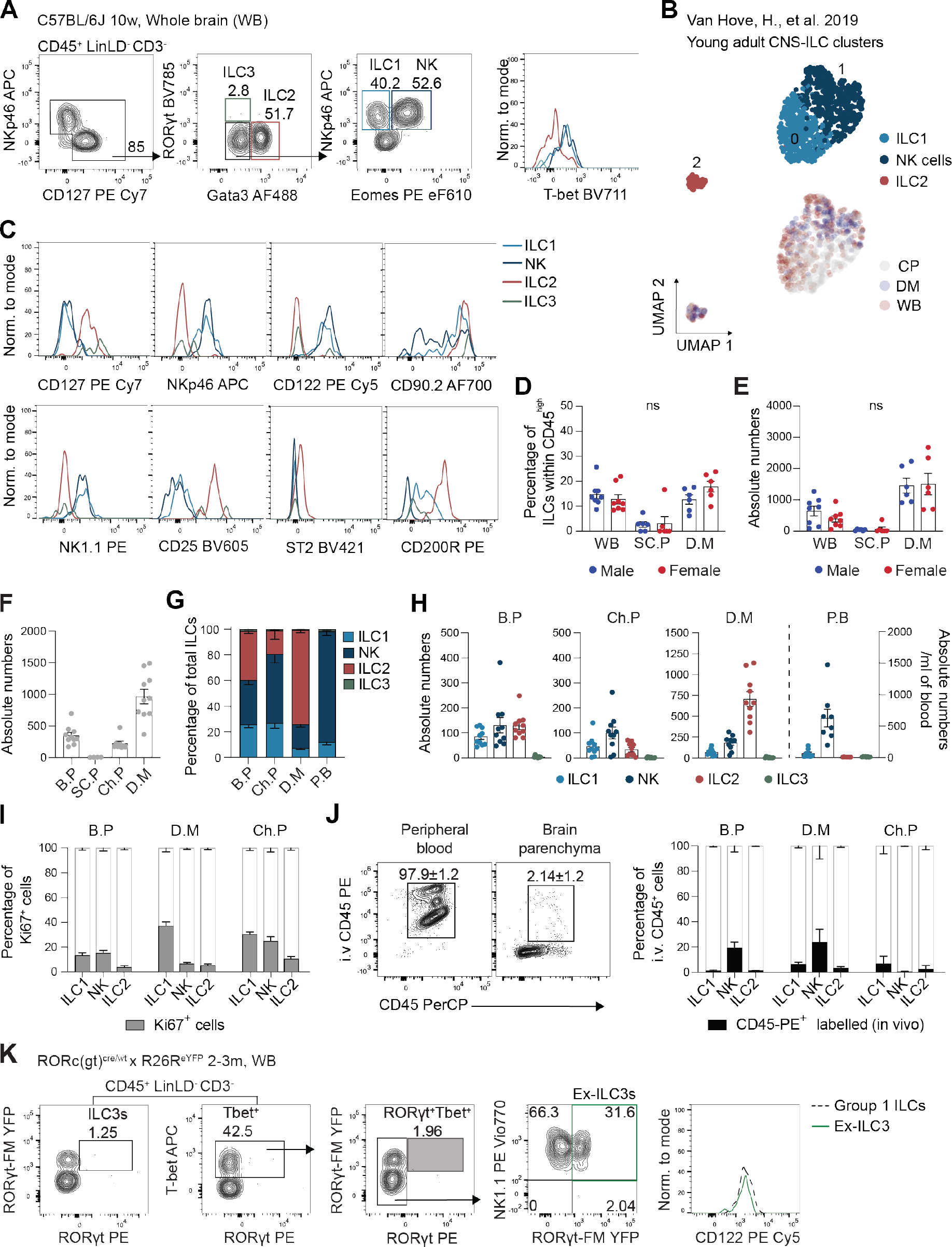
Young adult CNS harbors quiescent tissue-resident group 1 ILCs, ILC2s and ex-ILC3s. Flow cytometry analysis of young adult C57BL/6J (WT) 8-11 weeks old (8-10wo) mice. **(A)** Gating strategy to identify ILCs as CD45^high^LinLD^-^CD3^-^NKp46^+^ and/or CD127^+^ cells, whose subset identity is confirmed by the expression of the lineage-associated TF RORγt, Eomes, T-bet or Gata3, that defined ILC3s, group 1 ILCs and ILC2s, respectively. **(B)** Uniform Manifold Approximation and Projection (UMAP) visualization of scRNA-seq of ILC clusters found within the total CD45^+^ immune cells isolated from the whole brain or microdissected border regions (dura mater and choroid plexus) of 9-week-old C57BL/6 mice. Each colored cluster identifies a subset of ILC1-like, NK and ILC2s (above) or CNS-compartment of origin (below). Data are from Van Hove et al., (2019). **(C)** Representative histograms of markers expression by each identified ILC subset from young adult CNS. **(D)** Quantification of the percentage of total ILCs within CD45^high^ cells and their absolute numbers **(E)** in whole brain (with leptomeninges and choroid plexus), spinal cord parenchyma (SC.P) and dura mater (D.M) of male and female young adults (8-10wo). Date depicted as mean ± SEM., n = 8 examined over ≥ 3 independent experiments. D.M samples were pooled from two animals when necessary. **(F)** Absolute numbers of total ILCs in brain parenchyma (B.P, inc. leptomeninges), SC.P, choroid plexus (Ch.P) and D.M. Data depicted as mean ± SEM., n = 4- 10 examined over ≥3 independent experiments. D.M and Ch.P samples were pooled from two animals if needed. ILC subgroups **(G)** frequency and **(H)** absolute counts in B.P, Ch.P and D.M were compared to peripheral blood (P.B). Data are shown as mean ± SEM., n = 8-10 examined over ≥ 3 independent experiments. Dura meninges and choroid plexus samples were pooled from two animals when necessary. **(I)** Quantification of Ki67 expression among ILCs within the different CNS compartments. Data are depicted as mean ± SEM., n = 6. **(J**) Mice were intravenously injected with anti-CD45-PE (2 mg) 3 minutes before euthanasia and perfusion. On the left, the representative gating for identification i.v. CD45-PE^+^ immune cells (CD45-PerCP^+^) in peripheral blood and brain parenchyma with the quantification depicted as mean ± SD. The graph depicts the proportion of i.v. (intravascular) CD45^+^ cells among each ILC subset per compartment. Graph is depicted as mean ± SEM., n = 4 examined over 2 independent experiments. **(K)** Representative gating strategy for the identification of ILC3s and ex-ILC3s using the Rorc(gt)^cre/wt^×R26^eYFP^ fate map (FM)^+^ mice 2–3-month-old.

Confirming our and others’ previous work^6,33^, CNS group 1 ILCs, expressing NK1.1, NKp46 and CD122 (IL-2Rβ), conveyed not only conventional Tbet^+^Eomes^+^ NK cells but also Eomes^-^ ILC1s that were highly positive for CD90.2 and showed intermediate levels of CD200R expression (Fig. 1A, C; Sup. Fig. 1E). Mature ILC2s were identified by the expression of CD127, CD90.2, ST2 (IL-33R), and CD25 (IL-2Rα), together with high expression of CD200R (Fig. 1C). No significant differences were observed between female and male young CNS in terms of the proportion and absolute number of total ILCs or ILC subsets (Fig. 1D, E; Sup. Fig. 2A, B).

We found ILCs along all investigated CNS regions, i.e., brain parenchyma (including leptomeninges) (B.P, 348.2 ± 46.99), choroid plexus (Ch.P, 222.7 ± 36.8), and dura mater (D.M, 964.6 ± 114.9), but not in spinal cord parenchyma (SC.P, 3.32 ± 0.64), which was therefore excluded from subsequent experiments (Fig. 1E, F). In contrast to peripheral blood, ILC2s were the main population detected in the dura mater, whereas group 1 ILCs (both ILC1s and NK cells) were more prevalent in the brain parenchyma and choroid plexus (Fig. 1G, H). Strikingly, we observed negligible amounts of ILC3s in all CNS compartments (Fig. 1G, H).

CNS ILCs displayed a low proliferative capacity (Fig. 1I, Sup. Fig. 1F). Within the brain parenchyma, all ILC subsets showed proliferation rates below 20% (Fig. 1I) in both female and male animals (Sup. Fig. 2C), while a slightly higher proliferation rate was observed for group 1 ILCs in dura and choroid plexus (Fig. 1I). Furthermore, transcript analysis showed low levels of cytokine expression among the different ILC clusters, especially among ILC2s, further indicating low basal activity of ILCs in the steady-state CNS (Sup. Fig. 1F). Intravenous injections of anti-CD45-PE 3 min prior to euthanasia and perfusion, further demonstrated that the investigated populations excluded blood contaminant cells, identified as CD45-PE^+^ cells (Sup. Fig. 1G). Up to 90-99% of the tissue-resident helper-like ILCs were CD45-negative (Fig. 1J). In contrast, within the B.P and D.M, the NK cell fraction encompassed 25-35% of cells derived from the blood circulation or associated with the brain vasculature (Fig. 1J). In fact, scRNA-seq data also showed both; the helper-like clusters and the majority of NK cells presenting a tissue-resident signature (*Cd69, Rgs1 or Il1rl1* for ILC2s) and a fraction of NK cells showing higher expression of genes associated with a circulating phenotype (*Sell, S1pr1* and *Klf2*) (Sup. Fig. 1H). Helper-like ILCs were enriched in homing receptor *Cxcr6*, while *Ccr5* was predominantly present among NK cells and ILC1s. Interestingly, *Ccr2* was highly expressed among all ILC subsets (Sup. Fig. 1I).

Thus far, our data show that resting and quiescent tissue-resident ILCs are present within the adult CNS. While group 1 ILCs and ILC2s are enriched in the B.P and Ch.P, ILC2s are the main population in the D.M. Furthermore, although we had previously reported on the presence of ex-ILC3s within the steady-state CNS^6^, no ILC3s were observed along adult CNS structures. To complement our previous data, we used fate map *Rorc(gt)^cre^ × R26R^eYFP^* (RORγt-FM) mice and analyzed the presence of ex-ILC3s in 2-3-month-old mice. Using this reporter line, we confirmed the absence of ILC3s and the presence of ex- ILC3s principally among T-bet^+^ ILCs (Fig. 1K) and, in a much lesser extent, among T-bet^-^ST2^+^ ILCs (Sup. Fig. 1J) in adult mice. This data suggests an early presence of RORγt^+^ ILC3s within the CNS. T-bet^+^ ex-ILC3s expressed CD122 and NK1.1, and appeared predominantly within the whole brain (18.4 ± 3.5% within T-bet^+^ cells), while T-bet^-^ST2^+^ ex-ILC3s accounted for less than a 5% of the dura and brain ST2^+^ ILC2s (Sup. Fig. 1J).

### ILCs start infiltrating the brain prenatally as progenitor-like cells and mature ILCs

To investigate whether the presence of ex-ILC3s in the adult brain truly reflects the existence of RORγt^+^ ILC3s in earlier developmental stages, we focused on deciphering the dynamics of ILC seeding and niche establishment within the whole brain (excluding dura meninges) along ontogeny.

First, we investigated the presence of ILCs within the prenatal brain using Id2^GFP/+^ reporter mice, which is expressed in all ILCs. We could confirm that the expression pattern CD45^high^ Lineage^-^LD^-^CD3^-^CD122^+^ and/or CD127^+^ defines *bona fide* CNS Id2-GFP^+^ ILCs at the embryonic stage (Fig. 2A). Brain ILCs were nearly absent at E14 (Id2^+^CD122^+^: 4.43 ± 1.18%; Id2^+^CD127^+^: 2.16 ± 1.11%) but appeared clearly enriched at E16 (Id2^+^CD122^+^: 16.55 ± 5.39%; Id2^+^CD127^+^: 12.7 ± 5.24%) (Fig. 2B), positioning the beginning of the ILC seeding into the CNS between these two timepoints. Notably, we were able to identify CD122^+^CD127^-^ (P1), CD122^-^CD127^+^ (P3), and double-positive CD122^+^CD127^+^ (P2) cells (Fig. 2A), which lack CD5 and express other ILC markers such as CD90.2, CXCR3 (in the case of CD122-expressing cells) and CXCR6 (Fig. 2A, C). Although CD127^+^ cells (P2 and P3) express markers such as integrin α4β7 and c-Kit, they lack expression of CD135 (Flt3), excluding the presence of CLPs (live CD45^+^Lineage^-^CD3^-^CD5^-^CD127^+^α4β7^+/-^cKit^+^CD135^+^). Nevertheless, we identified PLZF^high^PD-1^+^ ILC progenitor-like cells among total *bona fide* CNS ILCs, especially enriched in the CD122^+^CD127^+^ (P2) population (ILC progenitor-like cells brain: 16.45 ± 11.7%) (Fig. 2D). These PLZF^high^PD-1^+^ ILC progenitor-like cells showed high expression of Gata3 (without expressing the ILC2-marker IL2rα (CD25), data not shown) and showed intermediate levels of RORγt and T-bet (Fig. 2E), which generally characterize ILC progenitors that are still able to differentiate into distinct ILC subsets^34^. In parallel, high and mutually exclusive expression of lineage-associated TFs (i.e. high T-bet-expression in CD127^-^CD122^+^ cells, enriched in group 1 ILCs) was also observed at this timepoint, indicating the co-existence of differentiated and committed ILCs among this first wave of embryonal ILC infiltration (Fig. 2E).

**Figure 2.**
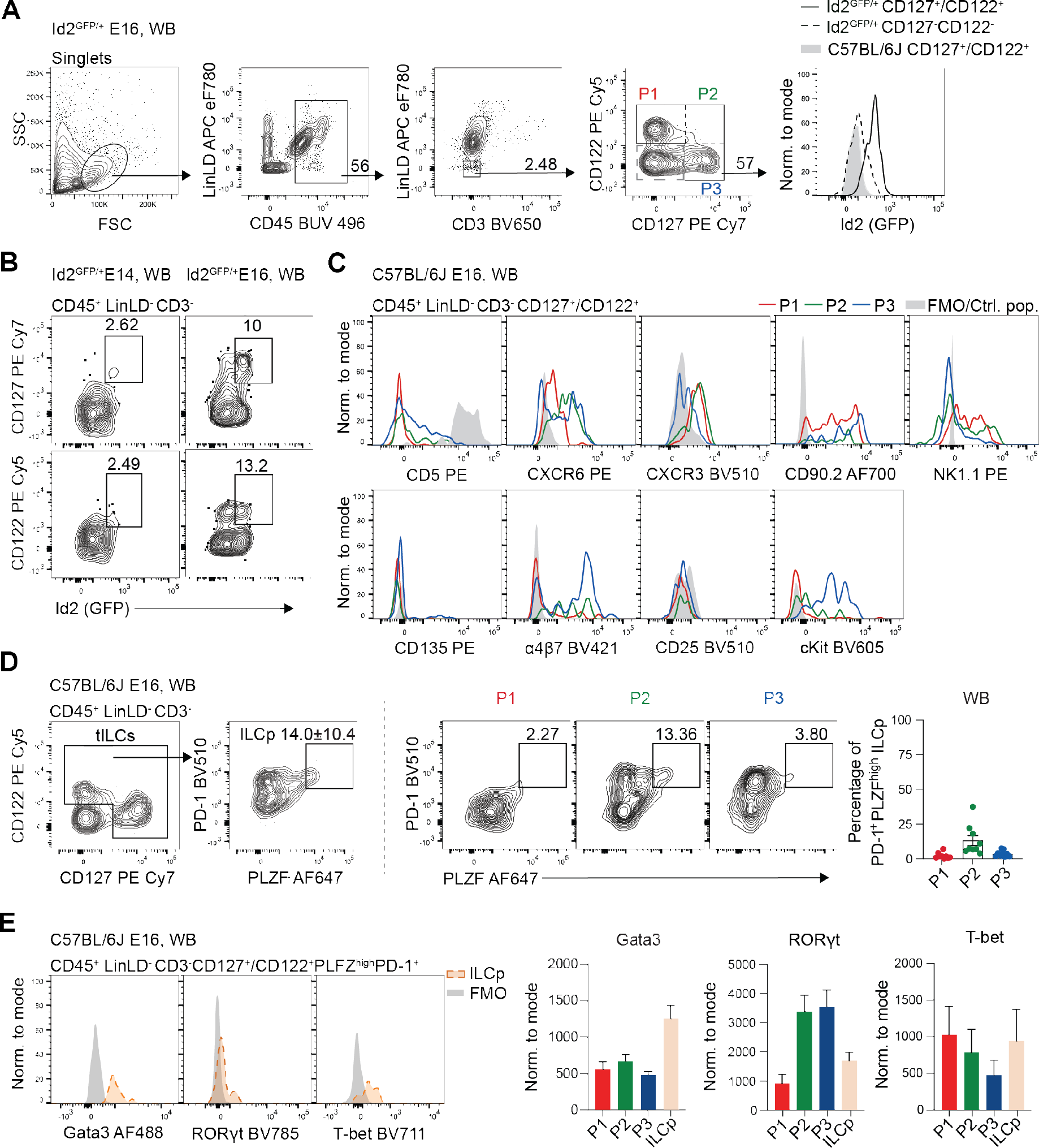
ILCs start infiltrating the brain prenatally as progenitors and committed ILCs. Flow cytometry analysis of E14 and E16 Id2^GFP/+^ reporter (indicated when used) and WT mice. Whole litters were pooled for each independent experiment. Analysis of Id2^GFP/+^ reporter animals includes WT and reporter animals within the pooled embryos. Of note, the rest of this study was always done in whole brain (inc. brain parenchyma, choroid plexus, leptomeningeal layers) and dura meninges was only analyzed separately when isolation was possible. **(A)** Representative gating of E16 Id2^GFP/+^ brain for identification of embryonic ILCs as CD45^high^LinLD^-^CD3^-^CD122^+^ and/or CD127^+^ cells whose identity is confirmed by the expression of Id2-GFP. **(B)** Quantification and representative flow cytometry of GFP^+^CD122^+^ and/or GFP^+^CD127^+^ ILCs from CD45^+^LinLD^-^CD3^-^ cells that start seeding the brain between E14 and E16 (n = 3 / timepoint). **(C)** Representative histograms of markers expression by CD122^+^CD127^-^ (P1), CD122^-^CD127^+^ (P3) and CD122^+^CD127^+^ (P2) populations from WT E16 brain (n = 2-6). Control population (Ctrl. pop.) used for CD5 defined as CD45^high^LinLD^-^CD3^+^. **(D)** Representative flow cytometry (left) of PD-1 and PLZF expression in CD45^high^LinLD^-^CD3^-^ CD122^+^ and/or CD127^+^ cells (total ILCs) and in P1, P2 and P3 (middle) from WT E16 brain. On the right, the quantification of these cells from the brain is depicted as mean ± SEM., n = 5-8 examined over > 3 independent experiments. **(E)** Representative histograms of TF expression measurement of PD-1^+^PLZF^+^CD122^+^CD127^+^ (ILCp) from WT E16 brains (left). Graphs (right) show the mean fluorescence intensity (MFI) ± SEM for each TF in ILCp compared to the rest of the cells in P1, P2 and P3 (n = 4-8). Data are representative of ≥ 3 independent experiments.

### Brain ILC seeding occurs in a subset-specific manner

Next, we investigated the seeding kinetics of lineage-committed ILCs during the pre- and postnatal stages starting at the identified timepoint of initial infiltration (E16). We defined CD45^high^LinLD^-^CD3^-^CD122^+^ and/or CD127^+^ cells as total ILCs and confirmed along ontogeny that the Id2^-^CD122^-^CD127^-^ population did not contain RORγt, T-bet, or Gata3-expressing cells (Fig. 3A). Interestingly, ILCs infiltrated the brain earlier than adaptive T lymphocytes (CD45^high^LinLD^-^CD3^+^), which were first detectable after birth (Fig. 3B). Furthermore, the calculation of ILCs per gram of brain showed that ILC numbers increased with brain weight, maintaining the ILC prevalence throughout the neonatal period and young adult life (Fig. 3C). Using RORγt, T-bet, or Gata3 to identify the different ILC subsets (Fig. 3A), we observed lineage-specific kinetics of brain seeding during development (Fig. 3D, E). CD122^+^T-bet^+^ group 1 ILCs was the largest population across all developmental timepoints, both in percentage and absolute numbers (Fig. 3D, E). Both Eomes^+^ and Eomes^-^ cells were identified pre- and postnatally (Sup. Fig. 3A). Until P1, Eomes^-^ cells appeared more predominant in frequency and absolute numbers (Sup. Fig. 3B, C), but their proportion decreased to 20-40% after the first week of life as the absolute numbers of Eomes^+^ cells increased (Sup. Fig. 3B, C). The increase in the Eomes^+^ population appeared to be caused by an influx of peripheral cells, as the proliferation rates of Eomes^-^ and Eomes^+^ cells were generally comparable (Sup. Fig. 3D). On the other hand, the second major population observed in the adult brain, Gata3^+^CD90.2^+^ ILC2s (ST2^+^, data not shown), appeared to start infiltrating the brain after birth, with increased prevalence and absolute numbers between P9 and 5w (Fig. 3D, E). Finally, we confirmed that RORγt-expressing cells were transitorily present perinatally, starting at very low levels at E16 but peaking by P9 (78.06 ± 21.9 cells/animal). Subsequently, their numbers started decreasing (P15: 35.2 ± 8.2 cells/animal; 5w: 53.07 ± 10.09 cells/animal) until almost complete retraction was observed in 10-week-old animals (Fig. 1A, G, H; Fig. 3D, E). Parallel analysis of peripheral blood ILCs along ontogeny showed differences in the prevalence of circulating ILC subtypes, indicating different waves of ILC mobilization (Fig. 3E, F). In this line, we observed a high proliferative capacity of all brain ILC subsets between E16 and P1 (Fig. 3G), which decreased progressively at later timepoints (Fig. 3G). Interestingly, as the proliferative capacity of embryonal CNS ILCs decreased at P9, the numbers in group 1 ILCs and ILC2s increased appreciably in the brain and peripheral blood circulation (Fig. 3E, F). Thus, we concluded here that a first wave of embryonal CNS ILC seeding combined with high levels of *in situ* proliferation drives the initial formation of the prenatal CNS ILC pool. However, the second wave of infiltrating ILCs during the first two weeks of life accounted for the further expansion of the brain ILC pool and the establishment of the ILC profile observed in the young adult CNS.

**Figure 3.**
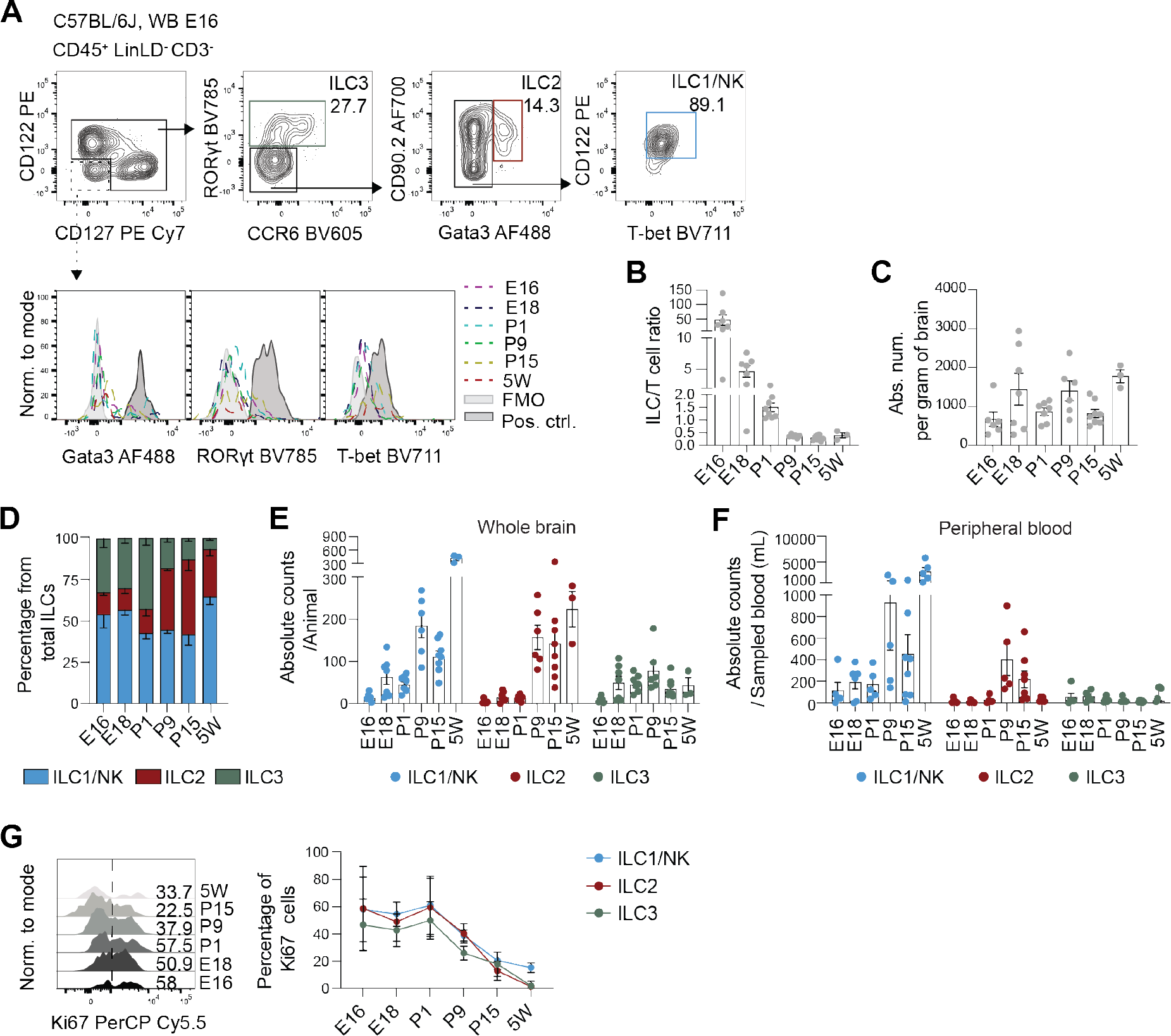
The CNS ILC pool formation is mediated by subset specific dynamics of tissue seeding. (A) Representative gating for the identification of the different ILC subsets in WT E16 whole brain: RORγt^+^ ILC3s, CD90.2^+^Gata3^+^ ILC2s and CD122^+^T-bet^+^ group 1 ILCs. The histogram representation along ontogeny of the CD122^-^CD127^-^ gate is depicted below. **(B)** ILC/T cell ratio over time in brain ontogeny (n = 3-7 / timepoint). **(C)** Absolute numbers of total ILCs (CD45^+^LinLD^-^CD3^-^CD127^+^ and/or CD122^+^) per gram of brain (n= 3-7/timepoint)**. (D)** Frequencies of group 1 ILCs, ILC2s and ILC3s along brain ontogeny (n = 3-7 / timepoint). **(E)** Absolute numbers of each ILC subset in whole brain and **(F)** peripheral blood (n = 6-7 / timepoint)**. (G)** Representative flow cytometry (left) and quantification (right) of Ki67 expression in each ILC subset in brain along ontogeny (n = 3-7 / timepoint). From E16 to P1, whole litter were pooled. From P9 to P15 onwards, 2-3 animals were pooled if needed. Blood was sampled from heart puncture with volumes: E16: ∼10 ul/animal, E18: ∼15 ul/animal, P1: ∼20 ul/animal, P9: ∼50 ul/animal, P15: ∼100 ul/animal, 5W: ∼300 ul/animal.

Altogether, we showed for the first time that ILCs start infiltrating the brain prenatally, earlier than adaptive T lymphocytes, in the form of both uncommitted PLZF^+^PD-1^+^ ILC progenitor- like cells and lineage-committed ILC1s, ILC2s, and, as hypothesized, ILC3s. Moreover, our data indicate that the formation of the CNS ILC pool is supported by a first wave of ILC infiltration followed by perinatal *in situ* proliferation and differentiation, whereas a second wave of infiltrating ILCs further expands the ILC pool after the first week of life.

### scRNA-seq analysis of postnatal ILCs reveals a mature ILC compartment with presence of group 3 ILCs

Analysis of RORγt-FM mice at different postnatal ages confirmed that ILC3s were present predominantly in the developing brain and peaked at P9 (Fig. 4A). To explore the properties of ILCs within the newborn CNS, we performed scRNA-seq analysis of CD45^+^LinLD^-^CD3^-^CD5^-^ CD122^+^/CD127^+^ ILCs from the whole brain and dura mater of P9.5 mice, as depicted in Figure 4B. This first-reported single-cell analysis of CNS ILCs was conducted with 480 cells from the brain and 362 cells from the dura mater after quality control and cell selection. We identified 5 distinct clusters of *Id2*-expressing cells with transcripts for *Gata3*, *Tbx21*, *Rorc,* and *Eomes,* as well as different associated ILC signature markers among their top differentially expressed genes (DEGs). ILC2s, cluster 0, expressing *Gata3, Il1rl1, Icos, Csf2*; ILC1s, cluster 1, expressing Tbx21, *Cd7, Ncr1, Nkg7, Il2rb,* and *Xcl1;* ILC3s, cluster 2, expressing *Rorc*, *Cxcr5, Cd4, Tcf7, Nrgn,* and *Lta*; and some NK cells, cluster 3, expressing *Tbx21, Eomes, Ccr5, Ccl5, Gmza,* and *Sell*. Cluster 4 represented a mixture of the previous clusters with additional markers for cell proliferation, such as *Mki67, Tuba1b,* and *Tubb5* (Fig. 4C, D; Sup. Fig 4A). Cells clustered based on ILC-subset specific identity markers and independently of the region of origin (Fig. 4C). Furthermore, the top 15 DEGs of each cluster were comparable between both investigated regions (Sup. Fig 4A). The *bona fide* ILC identity of the selected clusters was confirmed by the lack of (co)-expression of markers characteristic of other immune cell types of the CNS (Sup. Fig. 4B). No CLP or ILCp cells were identified, as demonstrated by the absence of a cluster of *Cd34* and *Flt3* or *Zbtb16* and *Tcf7* co-expressing cells, respectively (Sup. Fig. 4C). ScRNA-seq and flow cytometry analysis showed a similar region-specific subset distribution as the one observed in young adults, with group 1 ILCs as the main subset in the brain, closely followed by ILC2s and ILC3s. In the dura, ILC2s clearly appeared as the main ILC population (Fig. 4E).

**Figure 4.**
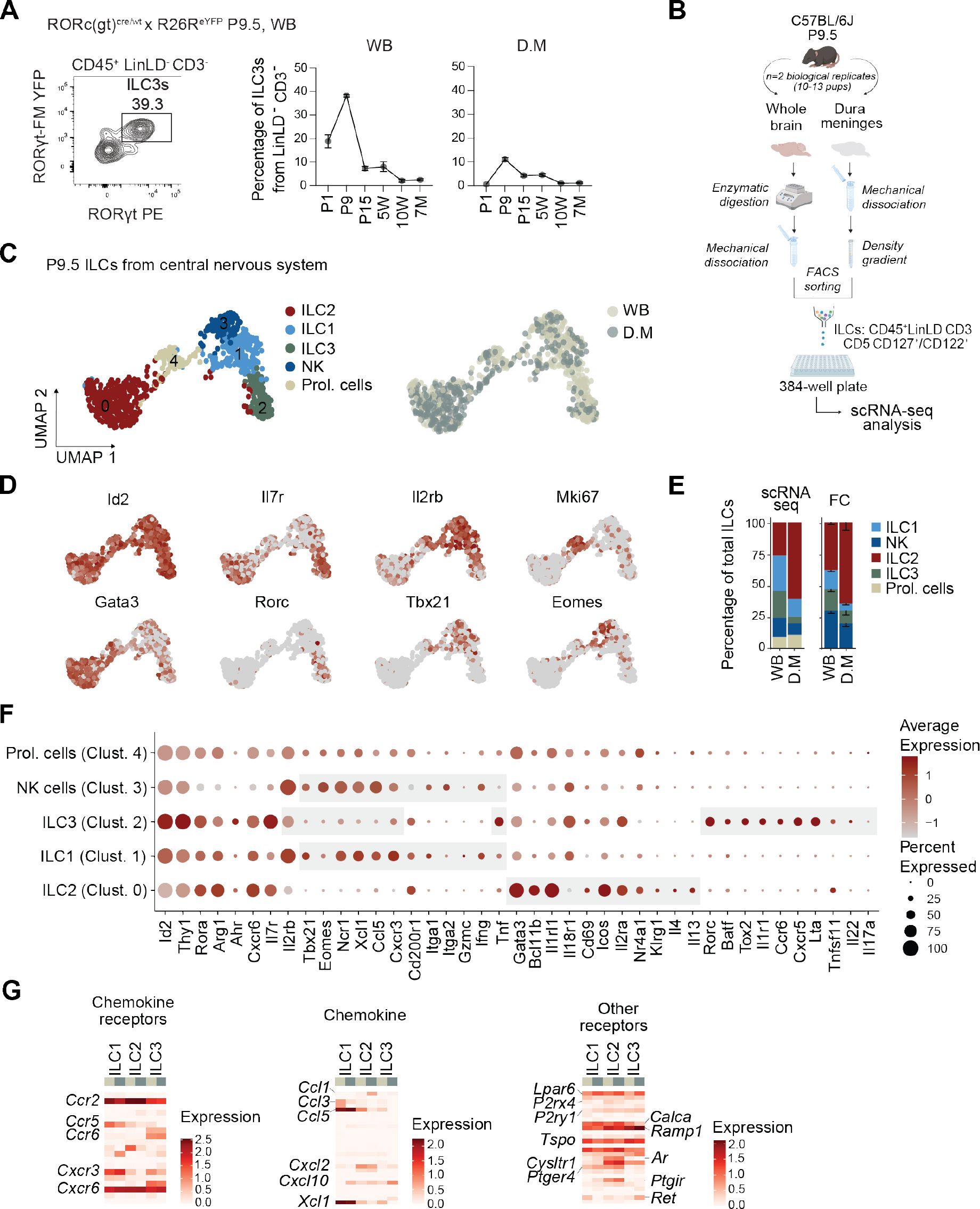
Single cell transcriptome profiling of newborn whole brain and dura cells shows a heterogeneous pool of mature ILCs. (A) Representative gating strategy for the identification of fate map (FM)^+^ ex-vivo RORγt^+^ ILC3s in P9 Rorc(gt)^cre/wt^×R26^eYFP^ mice (left) and quantification of YFP^+^RORγt^+^ ILC3s among CD45^+^LinLD^-^CD3^-^ cells depicted as mean ± SEM., n = 2-5 / timepoint from ≥ 2 independent experiments. At P1 FM^+^ pups from the litter were pooled. From P9 onwards animals were analyzed individually. **(B)** Schematic representation of the experimental set-up used to sort live CD45^+^LinLD (Fixable Viability Dye, CD19, B220, Cd11b, Cd11c, Gr-1, F4/80, FcεRIα)^-^CD3^-^CD5^-^ cells expressing CD127 and/or CD122 from brain and dura meninges, isolated from C57BL/6J P9.5 mice. Cells were single- cell sequenced by an adapted version of the SORT-seq protocol^67^. **(C)** UMAP depicted five distinct clusters along brain and dura meninges. **(D)** Gene expression UMAP plots of main subset-specific lineage markers. **(E)** Quantification of the frequency of ILC subsets identified by scRNA-seq and flow cytometry (FC) from total ILCs in brain and dura mater. For FC, percentages are shown as mean ± SEM., n = 4-6. **(F)** Dot plot depicting selected gene expression (average and percentage) within clusters. **(G)** Transcriptional analysis of helper- like ILCs (ILC1s, ILC2s and ILC3s), excluding NK and proliferating cells. Heatmap representing average gene expression levels of ILC-homing receptor, chemokines, neurotransmitter, and other receptors within ILC clusters in whole brain and dura mater.

We observed clear expression of signature ILC markers such as *Id2, Thy1, Rora, Arg1,* and *Cxcr6* among postnatal CNS ILCs (Fig. 4F). Brain and dura NK cells showed expression of *Eomes* and the integrin *Itga2* while ILC1s expressed higher levels of *Cd200r1* and *Itga1* (Fig. 4F). Furthermore, both populations of group 1 ILCs showed low *Tnf* but significant levels of *Ifng,* and *Ccl3* expression (Fig. 4F). Brain and dura ILC2s expressed similar levels of *Gata3, Icos*, and the IL25 and IL33 receptor subunits *Il1rl1* and *Il2ra* (Fig. 4F). Additionally, no or very low expression of *Il18r1* was detected together with expression of effector cytokines such as *Il4* and *Il13* (Fig. 4F), indicating predominant presence of more differentiated ILC2s. On the other hand, ILC3s were the population with higher expression of *Il7r* and *Thy1*, and molecules involved in ILC3 stability such as *Batf* and *Tox2* (Fig. 4F). Transcripts for both LTi (*Ccr6, Cxcr5, Lta* and *Tnfsf11*) and NKp46^+/-^ ILC3s (*Il2rb*, *Tbx21, Cd226, Cxcr3, Ccl5* and *Ncr1)* were observed (Fig. 4F). Of note, *Tnf*, but not *Il22* or *Il17f*, appeared as the main cytokine produced by CNS ILC3s at this developmental timepoint (Fig. 4F). Comparison of the expression of helper ILC-related chemokine, cytokines, and a plethora of cell receptors showed significant subset-specific but also compartment-specific ILC features. As demonstrated for adult ILCs (Sup. Fig. 1I) all CNS ILCs expressed high levels of the key mediators of migration *Ccr2* and *Cxcr6* (Fig. 4G). Of interest, ILC1s, especially within the brain, express Ccr5, Cxcr3 as well as Ccl3, Ccl4 and Ccl5 (Fig. 4G), and high levels of *Ifng* and *Tnf* (Sup. Fig. 4D). In contrast, ILC2s showed higher expression of different effector cytokines and chemokines such as Ccl1, Cxcl10, *Csf2*, *Il5,* and *Il13,* when located within the dura (Fig, 4G; Sup. Fig. 4D). The analysis of receptors involved in a possible response of CNS ILCs to neuropeptides, hormones, or metabolic signals (Sup. Table 2) indicated that, purinergic receptor (*P2rx4*), androgen receptor (*Ar*), prostaglandin receptors (*Ptger4, Ptgir*) or lipid receptor (*Cysltr1*) transcripts were enriched in ILC2s, again especially within the dura (Fig. 4G). This points to a higher ability of ILC2s to respond to neuroendocrine stimuli.

### Postnatal group 3 ILCs (incl. LTi and NKp46^+/-^ ILC3s) retraction results from ILC3 to ILC1 conversion and decreased cell turnover

To understand the mechanisms underlying ILC3 retraction from the developing brain, we performed hallmark pathway enrichment analysis based on the ILC3 cluster-specific genes (Fig. 5A). Interestingly, we observed highest activation of genes related to cell survival pathways such as the apoptosis and IL2/Stat5 signaling pathways (Fig. 5A). Further analysis showed that the ILC3 cluster appeared enriched in anti-apoptotic genes such as *Bcl2, Bcl2l1, Mcl1* and *Cflar,* in conjunction with key autophagy related genes (*Atg3, Atg7, Lamp1* or *Lamp2*) (Fig. 5B). This translated into a significantly higher ratio between the anti-apoptotic (Bcl-2) and the pro-apoptotic (Bim) molecules in ILC3s when compared to ILC1s (Fig. 5C), and low levels of ILC3 apoptosis measured by Annexin V and PI staining (data not shown). Of interest, although postnatal CNS ILCs progressively decrease their proliferative capacity (Fig. 3G), when assigning cell cycle scores based on the expression of S and G2M-associated genes, we identified not only a cluster of actively proliferating cells (cluster 4) but also a high proportion of cells in G2M phase distributed along the different ILC clusters (Fig. 5D). Cluster 4, enriched in proliferating cells, was found in similar proportion in brain and dura mater (Fig. 4E), and was enriched in group 1 ILCs and ILC2s (Fig. 5E, F). Flow cytometry analysis confirmed this data and showed ILC3s as the population with the lowest proliferation rate, especially brain ILC3s, which completely stop proliferating in later developmental timepoints (Fig. 3G; Fig. 5F). Overall, this indicated that ILC3s appeared to decrease their proliferation capacity while activating pro-survival mechanisms in the postnatal brain.

**Figure 5.**
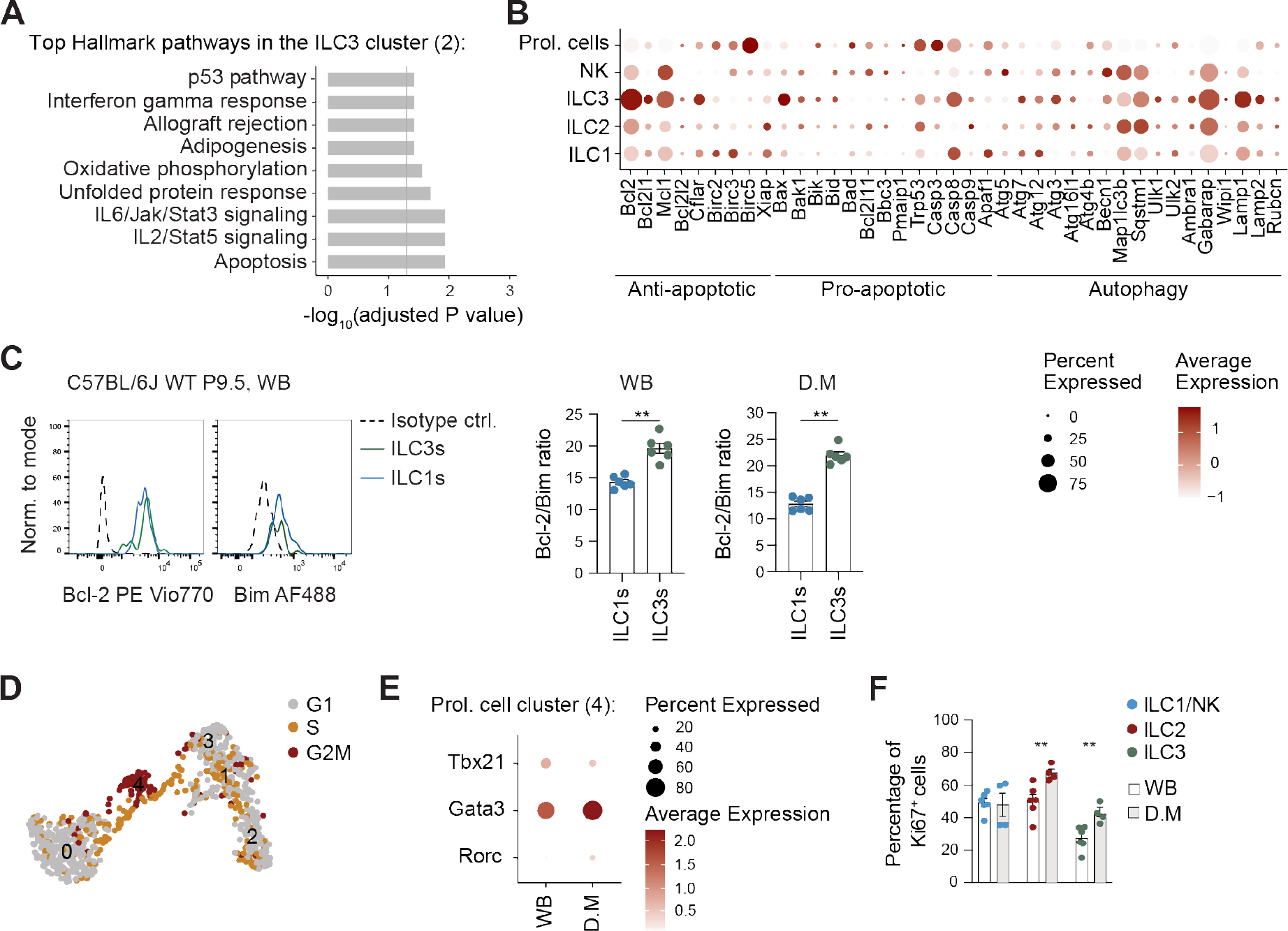
Postnatal ILC3s show high survival but low proliferation capacity. (A) Plot of the top pathways obtained by functional analysis on marker genes of cluster 2 (ILC3s) using the Hallmark pathway database. Bars represent the -log10 of the adjusted p values. **(B)** Dot plot depicting selected gene expression (average and percentage) of apoptosis and autophagy related markers within the identified clusters. **(C)** Representative flow cytometry plots of ILC1s (CD45^+^LinLD^−^CD3^-^NKp46^+^T-bet^+^Eomes^-^) and ILC3s (CD45^+^LinLD^−^CD3^-^ RORγt^+^). Histograms show expression of Bcl-2 and Bim in ILC1 (blue line) and ILC3s (green line) together with isotype controls (black dashed lines). Quantification of the expression ratio between Bcl-2 and Bim in the different populations. Ratios are shown as mean ± SEM, n = 6. Statistical test used is Mann-Whitney U test, **<0.01. **(D)** UMAP representation labeled by cell cycle phase. **(E)** Dot plot representation of the ILC-specific TFs among cluster 4 (proliferating cells). **(F)** Quantification by flow cytometry of the percentage of Ki67 expressing cells among ILC-subtypes in brain and dura mater. Percentages are shown as mean ± SEM., n = 4-6. Statistical test used is Mann-Whitney U test, **<0.01.

To further characterize the ILC3 population present in the developing CNS, we analyzed ILC3 subsets pre- and postnatally as well as in 5–week-old animals. As depicted in Fig. 6, CD4^+/-^ CCR6**^+^** LTi cells, NKp46**^+^** ILC3s and double-negative ILC3s were all present at perinatal stages (Fig. 6A; Sup. Fig. 5A). CD4^+/-^CCR6**^+^** LTi cells and double-negative ILC3s, appeared as the main populations until P9, whereas NKp46**^+^** ILC3s first appeared at P1 and peaked at P15 (Fig. 6A; Sup. Fig. 5A). Corroborating these results, unsupervised clustering conducted on the ILC1 and ILC3 superclusters of our scRNA-seq data from P9.5 mice, revealed 4 subclusters with top 15 DEGs: ILC1 (cluster 0: i.e. *Tbx21, Xcl1, Nkg7* and *Ifng*), Nkp46^+/-^ ILC3 (cluster 1; i.e. *Il7r, Rorc* and intermediate expression of *Il2rb, Ncr1,* and *Klrk1*), LTi (cluster 2; i.e. *Ccr6, Cd4, Cxcr5, Tnfsf11 and Il17f*) and a small cluster with low quality cells of unclear origin named as others (cluster 3; i.e. *Rgs4, Rgs7, Dmtn, Celf4, and Rtn1*) (Fig. 6B; Sup. Fig. 5B, C). Similar prevalence of the identified clusters was observed in brain and dura (Sup. Fig. 5D), although we clearly observed that ILC3s were predominantly located in the brain (and leptomeninges) (Fig. 4A; Fig. 4E). As previously shown, besides the changes in the prevalence of the different RORγt^+^ ILC subsets along development, a general contraction of this compartment was underlying the CNS maturation into adulthood (Fig. 3D, E; Fig. 4A; Sup. Fig 5A). We demonstrated a pronounced decrease in ILC3 turnover as the CNS developed (Fig. 3G, Fig. 4G-I), which was consistent among all ILC3 subtypes (Sup. Fig. 5E). No increased cell death was observed, however gene ontology pathway enrichment analysis based on the NKp46^+/-^ ILC3 and LTi subclusters-specific genes showed 1) high activation of pathways associated to active chemokine/cytokine signaling and leukocyte adhesion and, 2) upregulation of gene sets related to mononuclear/myeloid-leukocyte differentiation (Fig. 6C). To analyze possible plasticity events underlying this immune activation and differentiation, we performed pseudotime and trajectory analysis using Monocle 3 (cluster 1 as origin). A potential trajectory path was observed originating from NKp46^+/-^ ILC3 cluster and branching on one side towards the LTi cluster (2) and on the other side towards the ILC1 cluster (0) (Fig. 6D). Gradual decreased expression of ILC3-associated markers (*Tcf7, Tcrg-1, Tcrg-2, Ddc*), while increasing ILC1 markers such as *Cd52, Il2rb, Klrk1, Xcl1* or *Gimap4*, was observed along the NKp46^+/-^ ILC3 > ILC1 trajectory (clusters 1>0) (Fig. 6E). To confirm this assumption, brains of RORγt-FM mice were analyzed at different ages. A population of ex-ILC3s emerged at P9, 13.9 ± 1.2% (brain) and 8.04 ± 1.7% (dura mater), which was accompanied by a peak in the presence of double-positive RORγt^+^Tbet^+^ ILCs (Fig. 6F). Furthermore, although the double positive RORγt^+^Tbet^+^ intermediate population and ILC3s almost completely disappeared at later timepoints, the frequency of ex-ILC3s among Tbet^+^ ILC1s remained stable (between 10- 20%) throughout life (Fig. 6F; Fig. 4A). Overall, this indicated that RORγt^+^ ILCs retraction from the developing brain is also partially mediated by an *in situ* ILC3 to ILC1 conversion, probably, in response to changes in the tissue microenvironment.

**Figure 6.**
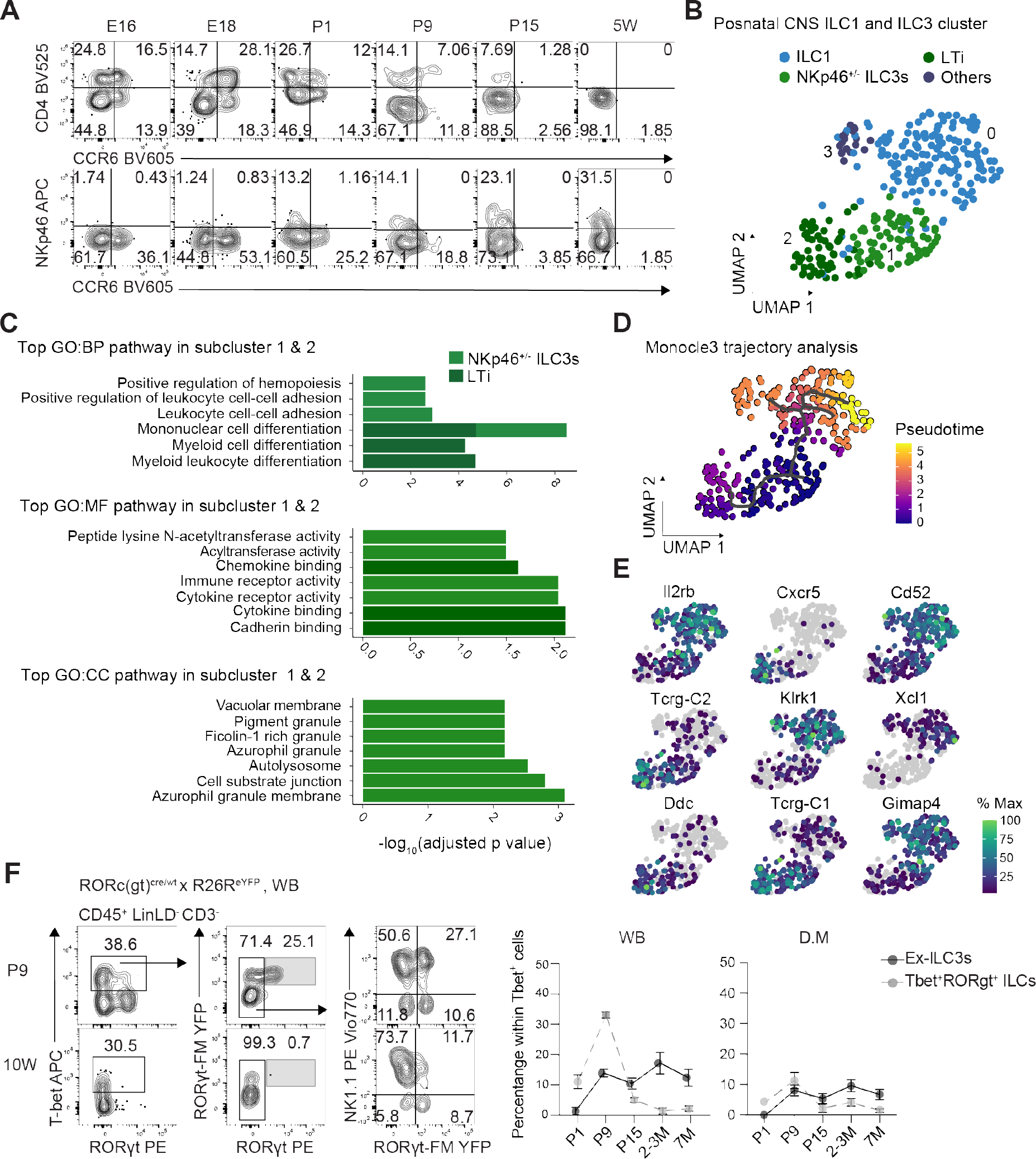
A heterogeneous pool of ILC3s is present perinatally and undergo ILC3>ILC1 conversion. (A) Representative plots of the expression of CD4, NKp46 or CCR6 by RORγt^+^ ILC3s along ontogeny. Mean frequencies are depicted in the graph. **(B)** UMAP dimensional reduction projection of ILC1/ILC3 superclusters reveals four separate subclusters. **(C)** Plot of the top pathways obtained by functional analysis on marker genes of cluster 1 (NKp46^+/-^ ILC3s) and cluster 2 (LTi) using the gene ontology biological process (BP), molecular function (MF) and cellular component (CC) databases. Bars represent the -log10 of the adjusted p values. **(D)** Trajectory analysis using Monocle 3 with origin cluster 1. Color-coded by cell distribution along pseudotime. **(E)** UMAP plots of the expression of the top genes driving the trajectory analysis projection. **(F)** Representative gating strategy for the identification of FM^+^ ex-ILC3 in P9 and 10W Rorc(gt)^cre/wt^×R26^eYFP^ mice (left) and quantification of YFP^+^ ex-ILC3s and double-positive RORgt^+^Tbet^+^ cells among Tbet^+^ ILCs depicted as mean ± SEM., n = 2-5 / timepoint from ≥ 2 independent experiments. From E16 to P1 whole litter were pooled. From P9 to P15 onwards, 2-3 animals were pooled if needed.

### Postnatal brain ILCs show an increased type 1 immune response

Next, to decipher possible local environmental features underlying the ILC3 to ILC1 conversion occurring predominantly within the brain, we compared the brain-specific ILC gene signatures with the dura as a reference compartment. Considering total ILCs, only 23 genes, appeared differentially expressed predominantly in brain by fold-change above or below 0.5/-0.5. The identified pattern was enriched in AP-1 components (*Junb, Fos, Fosb, Zfp36*) and GTPases of the immunity-associated proteins (GIMAPs*: Gimap3, Gimap4, Gimap5, Gimap6*), indicating higher immune activation and active modulation of cell survival/death pathways in brain ILCs (Fig. 7A). Gene set enrichment analysis (GSEA) of normalized counts showed an activated status of brain cells, characterized by the upregulation of gene ontology (GO) biological process (BP) pathways such as “response to calcium” and “activation of immune response”. In contrast, dura ILCs were enriched in metabolic-related pathways such as “small molecule metabolic process”, “ATP metabolic process” or “aerobic respiration” (Fig. 7B). The analysis of each brain ILC-subset separately revealed that for ILC1, ILC2, and ILC3, 189, 554, and 107 genes, were differentially expressed respectively, by fold-change above or below 0.8/-0.8 and Fisher p value < 0.05 (Fig. 7C). Gene set analysis (GSA) of the hallmark gene sets using the subset-specific identified DEGs and GSEA of normalized counts further confirmed that ILCs in the brain tended to express genes related to active immune response and differentiation (GSA-Hallmark: “allograft rejection”, “Tnfa signaling via NFKB”, “Interferon gamma and alpha response” and “reactive oxygen species pathway”); whereas ILCs in the dura showed higher activation of proliferative and metabolic-associated pathways (GSA-Hallmark: “Myc targets V1 and V2”, “oxidative phosphorylation” and “fatty acid metabolism”) (Fig. 7D). Interestingly, this active immune status within the brain was associated with type I signaling pathways (i.e. TNFa or interferon alpha/gamma immune response) (Fig. 7D).

**Figure 7.**
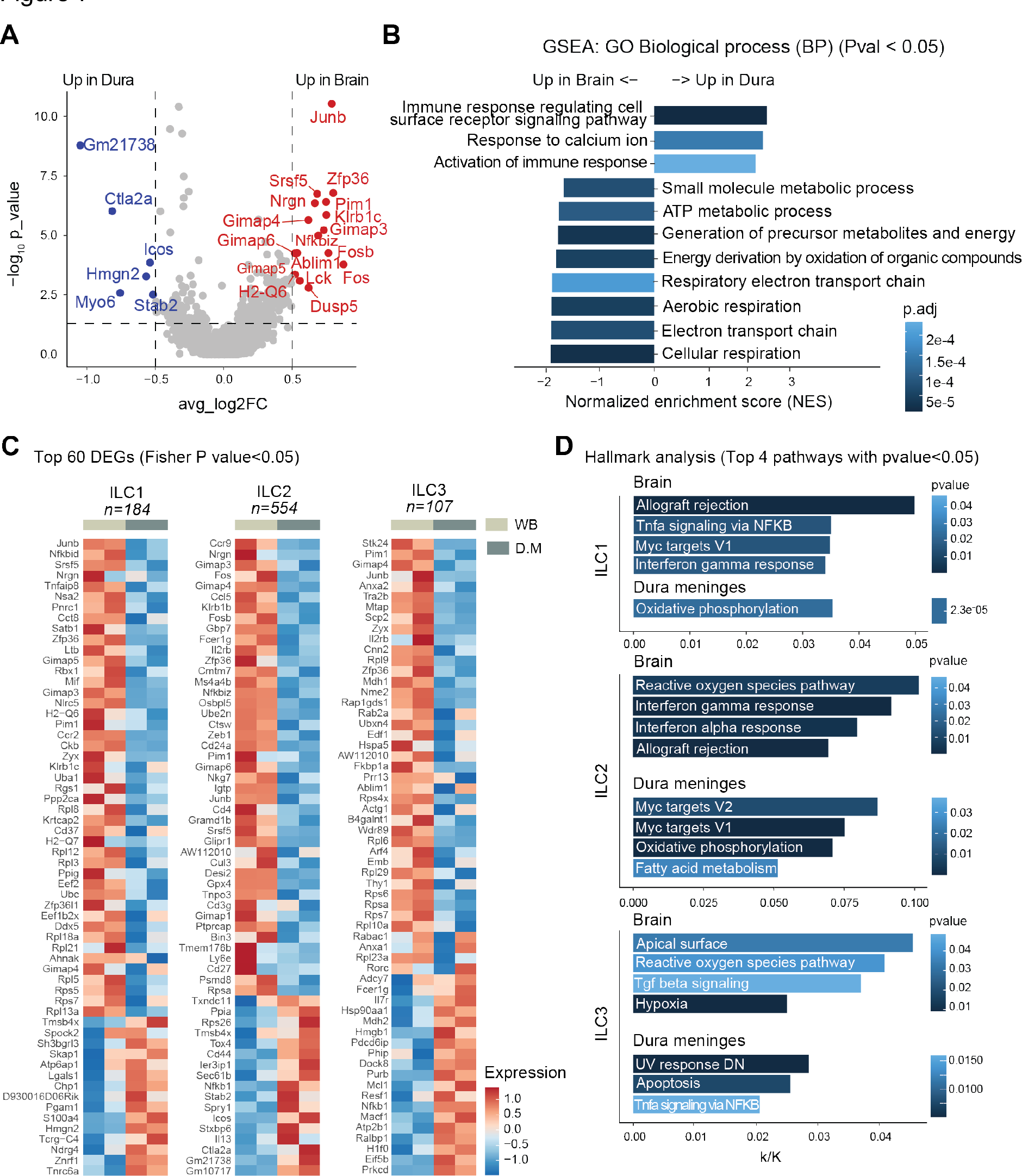
Regional differences are present between perinatal brain and dura ILCs. Transcriptional analysis of helper-like ILCs (ILC1s, ILC2s and ILC3s), excluding NK and proliferating cells. **(A)** Differential gene expression of ILC subsets in the brain and dura shown in a Volcano plot. Statistical analysis was performed by using non-parametric Wilcoxon rank sum. Red dots represent the genes highly expressed in brain, and blue dots represent the genes highly expressed in dura. **(B)** Results of GSEA GO biological process analysis showing enriched gene sets in brain vs. dura ILCs. The color in the bars indicate p adjusted values. A positive Normalized Enrichment Score (NES) value indicates enrichment in brain. A negative NES indicates enrichment in dura. **(C)** Heatmap of the DEGs for each ILC subtype in brain vs. dura selected based on Fisher p values < 0.05, resulted when performing Fisher exact P value test between single cell based non-parametric Wilcoxon rank test analysis and pseudobulk-based analysis using *lmFit, eBayes* and *topTable*. Expression values are represented as colors and range from red (high expression in brain), white (equal expression) and blue (high expression in dura). **(D)** Results of Hallmark analysis of identified differentially expressed genes in brain and the dura. The color depicts the p values meanwhile the bar size represents the k/K values of each pathway (ratio of number of DEGs divided by the number of genes in the indicated dataset).

**Figure 8.**
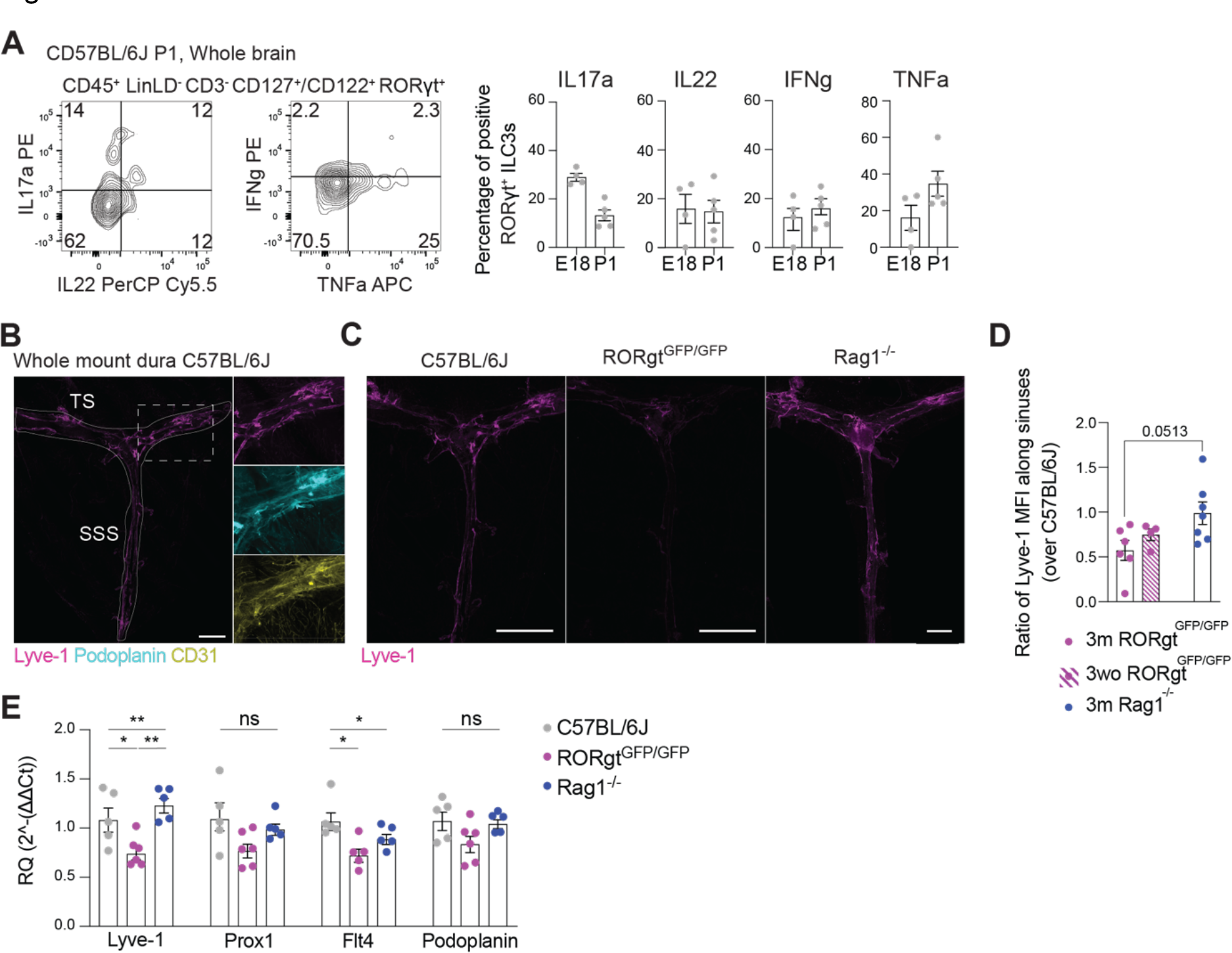
Altered lymphatic phenotype in the absence of RORgt^+^ ILCs but not Rag1^-/-^. **(A)** Representative plots (left) and quantification (right) of the expression of IL-17a, IL-22, INFg and TNFa among total brain RORγt^+^ ILC3s (n = 4-5) after stimulation with PMA/Ionomycin in the presence of Brefeldin A. Data are depicted as mean ± SEM. Data are representative of ≥ 3 independent experiments. Whole litter was always pooled for this analysis. **(B)** Representative images of Lyve-1-stained whole mount dura meninges showing lymphatic vessels located within the transversal sinus (TS) and the superior sagittal sinus (SS) (scale bar, 1000 µm). Square insets, higher magnification of Lyve-1, podoplanin and CD31 staining along the TS. **(C)** Representative images of control C57BL/6J, Rorgt-knockout (RORgt^GFP/GFP^) and T cell knockout (Rag1^-/-^) dura meninges collected and stained for lyve-1 (scale bar, 1000 µm). **(D)** Bar dot plots represent the ratio of the measurement of the Lyve-1 MFI staining along the TS and SSS between each knockout animal and its corresponding control C57BL/6J which was always analyzed in parallel. Ratios are shown as mean ± SEM., n = 4-7. Statistical test used is Mann-Whitney U test. **(E)** Expression of lymphatic-associated markers determined by RT-qPCR in the dura of C57BL/6J mice (controls), ILC3 deficient (RORgt^GFP/GFP^) and T cell deficient (Rag1^-/-^) mice. Relative quantification normalized to control animals is shown as mean ± SEM., n = 5-6. Statistical test used is Kruskal-Wallis test for three group comparison and Mann-Whitney U test for two group comparisons (* < 0.05; ** <0.01).

### Perinatal group 3 ILCs are functionally active and contribute to dura lymphatic development

To investigate the functional meaning of the transitory presence of ILC3, we assessed the production of IL-22, IL17a, TNFα and IFNγ of RORγt^+^ ILCs in perinatal stages E18 and P1, upon stimulation with Ionomycin/PMA. In contrast to P9 (Sup. Fig. 4D), we observed significant production of IL-22, but especially IL17a, among group 3 ILCs at earlier developmental timepoints (E18 and P1) (Fig. 7A). Interestingly, a peak in the production of the TNFα at P1, even before appearance of NKp46^+^ ILC3s, was detected (Fig. 7A). Among other signaling molecules, production of TNFα by ILC3s has been reported to be necessary for the formation of the gut associated lymphoid tissue^35^. Thus, we questioned whether the CNS RORγt^+^ ILCs could be involved in the formation of the lymphatic structures of the brain, i.e. the lymphatic vessels located within the sinuses of the dura meningeal layer, which have been reported to have the ability to acquire lymphoid-tissue like properties in the context of neuroimmune inflammation^36–38^. As shown in Fig. 7B, whole-mount dura meningeal expressed the endothelial lymphatic cell marker Lyve-1, podoplanin and showed intermediate expression of CD31 (Fig. 7B, zoomed images). In contrast, RORγt^GFP/GFP^ mice lacking RORγt^+^ ILCs and Th17 cells, but not T cell-deficient Rag1^-/-^ mice, showed a diminished expression of Lyve-1^+^ positive structures along the dural sinuses (Fig. 7C, D). This effect was observed both in adult mice (3m) and in 3-week-old mice (3wo), timepoint in which dura lymphatic vessels are reported to reach complete development^39^ (Fig. 7C, D). Expression analysis of lymphatic endothelial markers in the meninges of these mice also revealed a significant decrease of Lyve-1 and Flt4 expression, and a mild reduction of Prox1 and podoplanin, exclusively in RORγt^GFP/GFP^ animals (Fig. 7E). Altogether, our data indicates that RORγt^+^ ILCs, transiently present in the developing brain, play a role in the establishment of the lymphatic vessels of the dura meningeal layer.

## Discussion

The important role of the immune system in brain development and homeostasis has been revealed during the last decade^40–42^. Investigations on the implication of ILCs in such processes are however limited by their low prevalence within the healthy brain and the lack of knowledge regarding CNS ILC niche formation, maintenance, and functions. Using intravenous labeling and lineage-specific markers we describe here the full spectrum of *bona fide* resident ILCs in the healthy young adult CNS, which showed a compartment-specific distribution of differentiated cells. In line with previous studies^6,25,33^, we showed that group 1 ILCs, including ILC1 (NK1.1^+^NKp46^+^CD122^+^CD90.2^+^CD200R^int^Tbet^+^Eomes^-^) and NK cells (NK1.1^+^NKp46^+^ CD122^+^CD90.2^+/-^CD200R^-^Tbet^+^Eomes^+^), appeared predominantly located in inner brain structures (choroid plexus and brain parenchyma with leptomeninges), while ILC2s (CD127^+^CD90.2^+^CD25^+^ ST2^+^CD200R^high^Gata3^+^) were highly enriched in the dura meningeal layer. CNS ILCs of both, male and female young adult mice constituted a small population of mature lymphocytes that displayed a quiescent state with low basal levels of proliferative capacities and functional activation. This may suggest that in the normal developed brain compartments, innate lymphoid cells present low levels of basal activity and are able to promptly react to different stimuli, as reported by us and others in the context of autoimmune inflammation^6^, spinal cord injury^30^ or aging^26^. Interestingly, the presence of an ex-ILC3 population, principally among Tbet^+^RORγt^-^ ILC1s, pointed to dynamic changes in the CNS- ILC compartment during early life. To understand these changes, we have examined for the first time when and how the ILC compartment of the CNS is established and unraveled an important functional implication of perinatal ILC3s in CNS development and homeostasis.

The murine CNS is predominantly devoid of adaptive immune cells (T and B cells) at birth^40,41^. However, ILCs have been reported to start seeding peripheral tissues during fetal development in the form of progenitor cells and mature ILCs, appearing as early as E14.5 in tissues such as the intestine^10,11^ or the liver^15^. We therefore investigated ILC presence in the CNS (focusing on whole brain without dura meninges) pre- and postnatally using the Id2^GFP^- reporter line. Our data revealed a first appearance of Id2^+^CD45^+^LinLD^-^CD3^-^CD5^-^CD127^+/-^ CD122^+/-^ cells in the embryonal CNS, between E14 and E16. Although no CLPs were found within the CNS, we identified a small proportion of ILCs expressing high levels of α4β7, PD-1 and PLZF, and co-expressing intermediate levels of Gata3, T-bet and RORγt. This is compatible with a ILC progenitor-like identity, able to further differentiate into different ILC subsets^10^. The presence of this progenitor-like ILC population within the CNS supports the concept of brain seeding and *in situ* differentiation of ILC progenitor cells in non-hematopoietic organs during embryogenesis, as previously reported for intestine or lung^10,11^. Nevertheless, progenitor-like ILC cells seem to disappear from the CNS later in life as no cells with a progenitor identity were detected in our sequencing analysis at P9.5. We found, on the other hand, differentiated ILCs already at E16, indicating that the CNS ILC niche arises from both, *in situ* differentiation and expansion, as well as different waves of peripheral infiltration, mimicking mechanisms of layered ontogeny previously reported in peripheral organs^9,11,15,43^. In this line, we detected that brain embryonal Eomes^-^ and Eomes^+^ group 1 ILCs decreased their proliferation postnatally coinciding with a second wave of brain infiltrating Eomes^+^ cells. This second wave of infiltration occurred in parallel to the increase of Eomes^+^ group 1 ILCs in the peripheral circulation. In contrast, CNS ILC2s appeared to originate postnatally, as the main influx of ILC2s into the CNS occurs after the first week of life, concurrently with an increase in circulatory ILC2s. The dynamics of postnatal ILC2 infiltration coincides with those reported for other peripheral organs^9,44^. Finally, we showed a transitory presence of embryonal-derived ILC3s that disappear from the CNS after the first weeks of life. This ILC3 cell cluster seemed to mainly originate from embryonal ILCs that differentiate and/or expand within the perinatal brain as no significant numbers of circulating RORγt^+^ ILCs were observed at postnatal timepoints.

Altogether, we defined for the first time that the formation of the CNS ILC pool is established during the embryonal and postnatal life, since no further signs of infiltrating ILCs (besides a small fraction of NK cells) was observed in the healthy adult brain. Furthermore, we showed that group 1 ILCs and in lower levels, brain ILC2s and ILC3s, expressed type-1 associated chemokine receptors transcripts *Ccr5* and *Cxcr3*, whose ligands, CCL5 and CXCL9/10, have been reported to be expressed by brain-derived cells such as astrocytes^45^ or microglia^46^, respectively. This further suggests an implication of these chemokines in the perinatal iLC infiltration of the brain.

The postnatal ILC composition observed by flow cytometry was confirmed by scRNA-sequencing. We identified the complete spectrum of mature ILCs in newborn brain and dura matter, including group 1 ILCs (NK and ILC1s), ILC2s and ILC3s, with phenotypical and functional characteristics that resemble the ones obtained for peripheral tissues^10,47^. The compartment-specific separation observed in adult brains was already apparent in the newborn, with active group 1 ILCs enriched and proliferating in brain, while ILC2s were found more prevalent in the dura. This interesting compartmentalization also resembles that of other tissues, such as the lung or the liver, in which ILC2s are prominently localized at epithelial and fibroblast-rich barriers whereas type 1 lymphocytes accumulate within the parenchymal space^15,48^. In the lung, it has been reported that ILC1s control proliferation, infiltration and activation of ILC2s in an IFNγ-mediated manner^48,49^. Interestingly, our pathway analysis of the differentially expressed genes for each ILC subset in P9.5 showed an upregulation of genes related to IFNγ response especially in brain ILCs. Moreover, while ILC1 appears to be activated in this brain milieu, ILC2s and ILC3s displayed an increased expression of stress-response pathways, including reactive oxygen species production and hypoxia. Thus, the postnatal “inflammatory” parenchymal environment may favor survival and functionality of ILC1s, restraining other ILC subtypes, predominantly ILC2s, to meningeal barriers. Intriguingly, processes such as neurogenesis, synapse formation/pruning and blood-brain barrier formation, that may contribute to a state of controlled, focal tissue inflammation and lower nutrient and oxygen availability, are highly activated during the postnatal window in rodents^50,51^.

Focusing on the RORγt^+^ ILCs, the ILC3 pool shifted from being enriched prenatally with LTi cells to be composed principally by double negative and NKp46^+^ ILC3s after birth. Several processes appeared to be underlying the contraction of the RORγt^+^ population after the first week of life. First, in contrast to the basal levels of proliferation observed in group 1 and 2 ILCs, the ILC3 compartment seems to almost stop proliferating after birth, while upregulating expression of anti-apoptotic machinery mediators such as Bcl-2. Of note, Bcl-2 expression has been reported to fluctuate following changes in local cytokine signals such as IL-7^52^. Furthermore, we observed an increased expression of many autophagy-related genes (*Atg3, Atg7, Atg12, Lamp1* or *Lamp2*), suggesting ongoing processes of intracellular organelle degradation, which has been implicated in ILC survival^53^. This, together with the postnatal absence of peripheral blood circulating RORγt^+^ ILCs, translates into a drastic decrease of RORγt^+^ cell turnover within the postnatal brain. This resembles the ILC3 transitory presence also reported in the neonatal thymus, in which ILC3 also declined in numbers after birth while ILC2 becomes the main thymic ILC population^54^. In addition, analysis of RORγt-FM mice revealed the occurrence of *in situ* ILC3 to ILC1 conversion at postnatal day 9, a timepoint in which cells with a mixed phenotype were detected by both flow cytometry and single-cell sequencing analysis. The observed postnatal plasticity suggests that adult CNS ex-ILC3s are long-lived cells that originated during these first weeks of life as a reminiscence of the perinatal presence of RORγt^+^ ILCs. Indeed, presence of embryonal derived ILCs have been reported also in other adult tissues^9,12,15^, proving the ability of these cells to remain as long-life cell within the tissue. Finally, the activation of inflammation-related and cytokine signaling pathways (type 1 immune response) that seems to occur within the postnatal brain could also contribute to the contraction of ILC3s from the brain and the concurrent ILC3 to ILC1 conversion as reported in peripheral tissues in both, human^55^ and mice^20,56,57^.

On the functional level, we observed high levels of TNFα production by ILC3s at pre- and postnatal timepoints. RORγt^+^ lymphocytes associated signaling molecules such as TNF and LTα seem to be necessary for the development of secondary lymphoid structures, including lymph nodes but also the intestinal associated lymphoid tissue, Peyer’s patches (PPs)^35,58,59^. In line with this, we showed here that the absence of RORγt^+^ ILCs in the developing brain affected the formation of the newly described lymphatic vessels of the dura mater^60^. This was already observed at P15, timepoint in which the dura lymphatic vessel formation should be completed^39^. It has been shown that vascular endothelial growth factor C (VEGF-C)/VEGF receptor 3 (VEGFR3) signaling is essential for the formation of these lymphatic vessels^61^. However, TNFα appears to also cooperate with this axis in the promotion of lymphangiogenesis, through indirect activation of tissue macrophages during pathological conditions such as cancer^62^. Due to the unique timeline of dura lymphatic formation, we could speculate that TNFα- production by perinatal ILC3s contribute to the process of dura lymphatic formation, revealing an unique involvement of RORγt^+^ ILCs-derived signaling in dural lymphangiogenesis.

## Methods

### Mice

Rorct^m2litt^/J mice (Eberl et al., (2004))^63^, used as Rorc(gt)^GFP/GFP^ and Rorc(gt)^GFP/+^ mice, Id2^GFP/+^ mice (Rawlins et al., (2009))^64^ and Rag1^-/-^ mice (Mombaerts et al., (1992))^65^ were provided by Andreas Diefenbach (Charité-Universitätsmedizin Berlin, Berlin) and bred in our animal facility. Rorc(gt)-Cre^Tg^ (Eberl et al., (2004))^63^ mice crossed to Rosa26RE^eYFP^ mice (Srinivas et al., (2001))^66^, used as Rorc(gt)-fate mapping animals, were provided by Chiara Romagnani (DRFZ, Berlin) and bred in our animal facility. Rorc(gt)-cre was carried only by female breeders to prevent germline YFP expression. Non-transgenic experiments were performed with female and male, wild-type C57BL/6J mice purchased from the Research Institute for Experimental Medicine (FEM) of the Charité (Berlin, Germany) and Charles River Laboratories (Freiburg, Germany). All mice used were between 8 to 16 weeks old, unless indicated otherwise. Animal were pooled, when necessary, as indicated along the text.

All mice were bred under specific pathogen-free conditions in the Research Institute for Experimental Medicine of the Charité (Berlin, Germany). Animal handling and experiments were conducted according to the German and European animal protection laws and approved by the responsible governmental and local authority (Landesamt für Gesundheit und Soziales).

### Organ collection

Peripheral blood samples were collected in tubes with 2 mM EDTA at room temperature from euthanized mice before perfusion with 1-5 mL (embryos), 5-10 mL (pups) and 40-50 ml (adults) of ice-cold PBS. The rest of the organs: brain and spinal cord (with pia matter), choroid plexus, dura meninges (including partially arachnoid meninges), spleen, lymph nodes and liver were collected on ice-cold medium/PBS and processed immediately. For RT-qPCR, samples were lysed in RLT buffer (lysis buffer) and stored at -20°C. For immunofluorescence, transcardial perfusion with ice-cold PBS was followed by perfusion with 10-20 mL of buffered 4% paraformaldehyde (PFA, EMS).

### Tissue processing

The CNS (brain and/or spinal cord) was mechanically homogenized and passed through a 70 μm cell strainer (Corning) with complete medium [RPMI-1640 supplemented with 2 mM L-glutamine (Gibco), 100 U/mL penicillin (Seromed), 100 μg/mL streptomycin (Seromed), 10% FCS (Sigma–Aldrich) and 1% HEPES (Gibco)]. After centrifugation at 400 g for 15 min at 4°C, the pellet was resuspended in 5mL of 40% Percoll (GE Healthcare) and the lymphocytes were collected from the pellet after centrifugation at 2,200 g. Erythrocyte lysis was performed when necessary for 5 min at room temperature in between the two following washing steps done with PBS/BSA.

For the analysis of the choroid plexus separately from the brain parenchyma, the tissue was removed under a dissecting microscope from the lateral, third and fourth brain ventricles, and processed in parallel. Brain parenchyma processing was performed as described above. Isolated choroid plexuses were digested in 500 μL PBS containing 1 mg/ml Collagenase/Dispase (Sigma) and 1 mg/mL DNAse I (Sigma) for 20 min at 37°C with 400 rpm. In parallel, the dura meninges were peeled off from the interior side of the skull cap and digested in 500 μL PBS containing 2.5 mg/ml collagenase type VIII from Clostridium Histolyticum (Sigma) and 1 mg/ml DNAse I (Sigma) for 20 min at 37°C with 400 rpm. After digestion, both tissues were passed through a 70 μm cell strainer (Corning) and centrifuge at 400 g for 10 min.

Single cell suspensions of spleen, lymph nodes and liver were obtained by homogenizing the tissue through a 100 μm cell strainer (Corning). The spleen and blood underwent erythrocyte lysis for 10 min and washed one or two times as needed. Liver lymphocytes were enriched after mechanical dissociation and homogenization through a 100 μm cell strainer by centrifuging at 50 g for 1 min and discard the pellet.

### Labelling of the vascular compartment

To distinguish vasculature-associated circulating cells from those residing within the CNS, 2 ug/200 ul of PE anti-mouse CD45.2 antibody (Biolegend, 30-F11) diluted in PBS solution were injected through the tail vein. Animals were sacrificed 3 min post-injection.

### Flow cytometry

Extracellular flow cytometry staining was performed at 4°C in PBS containing 0.5% BSA. Dead cells were excluded by staining with Fixable Viability Dye (LD) (eBioscience, 1:4,000). Then, Fc receptors were blocked by incubating 15 min with anti-mouse CD16/CD32 (clone 2.4G2, BD Biosciences). The following antibodies were used for staining: PerCP anti-mouse CD45 (Biolegend, 30-F11, 1:100) or BUV496 anti-mouse CD45 (BD, 30-F11, 1:100), APC-Cy7 anti mouse/human CD45R/B220 (Biolegend, RA3-6B2, 1:100), APC-eFluor 780 anti-mouse CD45R/B220 (eBioscience, RA3-6B2, 1:100), APC-eFluor 780 anti-mouse CD19 (eBioscience, eBio1D3, 1:100-1:200), APC-Cy7 anti mouse-CD11b (BD, M1/70, 1:50), APC-eFluor 780 anti-mouse CD11b (eBioscience, M1/70, 1:50), APC-eFluor 780 anti-mouse Gr-1 (eBioscience, RB6-8C5, 1:100), APC-eFluor 780 anti-mouse FcεRI (eBioscience, MAR-1, 1:200), APC-Cy7 anti mouse F4/80 (Biolegend, BM8, 1:100), APC-eFluor 780 anti-mouse F4/80 (eBioscience, BM8, 1:100), APC-eFluor 780 anti-mouse CD11c (eBioscience, N418, 1:100), BV650 (Biolegend, 17A2, 1:50) or PerCP-Vio 700 (Miltenyi, REA641, 1:200) anti- mouse CD3, BV510 anti-mouse CD4 (Biolegend, RM4-5, 1:50), PE anti-mouse CD5 (Biolegend, 53-7.3, 1:300), PE-Cy7 (Biolegend, 1:50), PE (Biolegend, 1:100) or BV421 (Biolegend, 1:50) anti-mouse CD127 (A7R34), biotin (Biolegend, 1:200) or PE-Cy5 (Biolegend, 1:300) anti-mouse CD122 (TM-β1), Streptavidin-PE-Cy5 (1:300), PE anti-mouse CD200R (Biolegend, OX110, 1:200), Alexa Fluor 700 anti-mouse CD90.2 (Biolegend, 30- H12, 1:100), APC (Biolegend, 1:50), BV711 (Invitrogen, 1:50) or BUV395 (BD Horizon, 29A1.4, 1:50) anti-mouse NKp46 (29A1.4), BV421 anti-mouse α4β7 (BD, DATK32, 1:100),

BV421 (Biolegend, 1:50) and PerCP Cy5.5 (Biolegend, 1:50) anti-mouse ST2 (DIH9), BV605 (Biolegend, 1:50) and BV510 (Biolegend, 1:50) anti-mouse CD25 (PC61), PE-Vio615 anti- mouse CXCR5 (Miltenyi, REA215, 1:50), PE anti-mouse CD135 (Biolegend, A2F10, 1:100), BV605 anti-mouse c-Kit (Biolegend, ACK2, 1:100), BV510 anti-mouse CXCR3 (Biolegend, CXCR3-173, 1:50), PE anti-mouse CXCR6 (eBioscience, DANID2, 1:50), BV510 anti-mouse PD-1 (BioLegend, 29F.1A12, 1:100), BV605 anti-mouse CD196 (CCR6) (Biolegend, 29-2L17, 1:200) and PE and PE-Vio770 anti-mouse NK1.1 (Miltenyi, REA1162, 1:50).

For intracellular and intranuclear staining, the FoxP3 transcription factor staining buffer set (Invitrogen) was used to fix and permeabilize the cells according to the manufacturer’s instructions. In case of reporter or fate-map GFP/YFP signal, cells were fixed for 20 min at room temperature with 2% PFA (EMS) before beginning with the FoxP3 staining protocol. Cells were stained with BV711 (1:50) or AlexaFluor647 (1:300) anti-mouse T-bet (Biolegend, 4B10), PE eFluor610 anti-mouse Eomes (Biolegend, Dan11mag, 1:200), AlexaFluor488 (1:50) or AlexaFluor647 (1:50) anti-mouse Gata3 (Biolegend, 16E10A23), PE (BD, 1:500) or BV786 (BD, 1:100), anti-mouse RORγt (Q31-378), AlexaFluor647 anti-mouse/human PLZF (BD, R17-809, 1:800), PerCP-eFluor 710 anti-mouse Ki67 (eBioscience, SolA15, 1:1000), AlexaFluor488 anti-mouse Bim (Cell Signaling, C34C5, 1:100), AlexaFluor488 rabbit anti- mouse mAb IgG XP Isotype Control (Cell Signaling, DA1E, 1:100), PE-Vio770 anti-mouse Bcl-2 (Miltenyi, REA356, 1:100) and PE-Vio770 human anti-mouse IgG1 (Miltenyi, REA294, 1:100). Apoptosis analysis was done with the APC Annexin V (1:50) apoptosis detection kit with PI (1:200) in Annexin V Binding Buffer (Biolegend).

For the analysis of cytokine production, CNS derived single-cell suspension were stimulated in 96 well plates in complete medium with PMA (10ng/ml, Sigma-Aldrich) and Ionomycin (500ng/ml, Sigma-Aldrich), and cell transport inhibitor Brefeldin A (10 μg/ml, Biolegend) was added 1 hour later for a total incubation time of 4 hours at 37°C. After washing, cells were extracellularly stained and fixed with 2% PFA (EMS) for 20 min at room temperature. Once cells were fixed, the FoxP3 staining kit was used again to do a second fixation and permeabilization. Cells were then stained with PE anti-mouse IFNγ (Biolegend, XMG1.2, 1:100), APC anti-mouse TNFα (BD Bioscience, MP6-XT22, 1:50), PE anti-mouse IL-17A (Biolegend, TC11-18H10.1, 1:200) and PerCP Cy5.5 anti-mouse IL-22 (Biolegend, Poly5164, 1:25) for 45 min at room temperature.

Sample acquisition was performed using a LSR Fortessa flow cytometer (BD Biosciences). Data were further analyzed in FlowJo Software v.10 (FlowJo). Gating of populations were defined with fluorescence minus one (FMO) staining controls when necessary. Flow cytometry plots are shown as contour plots (5% with outliers). Overlay histograms were normalized to mode.

### Immunofluorescence staining and image analysis

Transcardially perfused mice were decapitated immediately and the skull cap was removed and clean of overlying skin and muscle before being drop fixed in 2% PFA at 4°C for 24 hours and washed with PBS for another 24 hours. The dura/arachnoid was then carefully peeled from the skull cap using fine surgical forceps and stored in PBS at 4°C. Whole mounts were stained in 24-well plates with constant agitation following the same protocol as in Louveau et al., (2015)^60^. WT animals and mutant were always stained in parallel for paired comparison. First, they were blocked and permeabilized with PBS containing 2% horse serum, 1% BSA, 0.1% Triton-X-100 and 0.05% of Tween 20 for 1 hour at room temperature, followed by incubation with primary antibodies: purified rat anti-mouse CD31 (BD Pharmingen™, MEC 13.3, 1:100) overnight at 4°C in PBS containing 1% BSA and 0.5% Triton-X-100. Whole- mounts were then washed 3 times for 5 min at room temperature in PBS followed by incubation with Alexa Fluor 568 anti-rat (Invitrogen, 1:500) for 1 hour at room temperature in PBS with 1% BSA and 0.5% Triton-X-100. Meninges were washed 3 times again for 5 min at room temperature in PBS and incubated with eFluor 660 anti-mouse Lyve-1 (eBioscience, ALY7, 1:200) and AlexaFluor488 anti-mouse Podoplanin (eBioscience, eBio8.1.1 (8.1.1), 1:200). Finally, nuclei were stained with 1:10,000 DAPI reagent, mounts were washed with PBS and mounted with Epredia™ Shandon™ Immu-Mount™ under coverslips.

Images were acquired with a Nikon Scanning Confocal A1Rsi+ and/or a Keyence BZ-X810 fluorescence microscope. Image analysis was performed using FIJI plugin for ImageJ (National Institutes of Health). Analysis of meningeal Lyve1 was performed measuring the mean fluorescence intensity of Lyve1 of the whole dura meninges along 10 mm of Z-stack images taken every 1 mm. The mean of the values obtained per plane was calculated and the ratio between the mean fluorescence intensity of mutant versus its age-and gender matched WT control was calculated.

### Real-time quantitative PCR (RT-qPCR)

mRNA was isolated from dura meninges by using RNeasy Mini Kit (Qiagen) following the manufactureŕs protocol. RNA samples were further transcribed into first strand complementary DNA by using High-Capacity cDNA Reverse Transcription kit (Applied Biosystems). RT-qPCR reactions were assayed in triplicates per sample using a QuantStudio^TM^ 5 Real-Time PCR system and TaqMan Gene expression assays (all Thermofisher): Lyve-1 (Mm00475056_m1), Podoplanin (Mm01348912_g1), Flt4 (Mm01292604_m1), Prox1 (Mm00435969_m1) and Hprt (Mm03024075_m1). mRNA content was normalized relative to the mean expression of the arithmetic means of Hprt CT values by applying the comparative CT method (2−ΔCT) in which ΔCT (gene of interest) = CT (gene of interest) − CT (arithmethic mean of housekeeping reference value). Furthermore, the ratio was calculated based on C57BL/6J control samples.

### ILCs scRNA sequencing and analysis

Brain and dura meninges were collected and processed as describe in “Tissue processing”. Immune cells from P9.5 C57BL/6J mice were collected in two biological replicates with a total of 10-13 pups per replicate. Cells were sorted as CD45^+^LinLD^-^CD3^-^CD5^-^CD127^+^ and/or CD122^+^into 384-well cell-capture plates containing barcoded primers and mineral oil and, immediately stored at -80°C. Plates were processed and sequenced at Single Cell Discoveries (Utrecht, The Netherlands), using an adapted version of the SORT-seq protocol^67^ with primers described in Van den Brink et al., (2017)^68^. Library preparation was done following the CEL- Seq2 protocol^69^ to prepare a cDNA library for sequencing using TruSeq Small RNA primers (Illumina). The DNA library was paired-end sequenced on an Illumina NextSeq 500, high output, with a 1 × 75 bp Illumina kit. Reads were mapped on Mus musculus (GRCm38) using STARsolo (2.7.10a). The percentage of retained wells range between 48%-79% with 44.45 to 55.22 million retained reads per library. Both run results were combined and cells were filtered based on the expression of 500-4000 genes with mitochondrial counts below 10%, leaving 480 cells from brain and 362 from dura meninges for subsequent analysis. Seurat 5.0.1 was used for further analysis.

### Dimensional reduction and clustering

Using functions of the Seurat package (v5.0.2), we first perform standard pre-processing (log-normalization with the *NormalizeData()* function, scale factor=10000) and identify variable features based on variance stabilization transformation for which the top 2000 highly variable genes were selected (*FindVariableFeatures()* function, method “vst”). Next, we scaled the integrated data using the *ScaleData()* function (“nCount” and “mitochondrial percentage” were used for regression). Once differences generated from technical preparations were minimized, principal component analyses (50 principal components) were performed on variable genes and embedded in 2-dimensional UMAP plots. We used jackstraw test to calculate the statistical significance and variance of each principal component. Based on these results, we performed clustering using the *FindNeighbors()* and *FindClusters()* methods with the first 12 principal components and a resolution of 0.7. ILC identity was confirmed by discarding co-expression of other immune cells specific genes (see Sup. Fig. 4B) and by confirming expression of canonical ILC genes among the top DEGs of each cluster and assessing expression of known ILC subset marker genes along the clusters (Fig. 4D, F and Sup. Fig. 4A).

### Differential expression

Compartment (brain or dura) specific markers were then identified using the Wilcoxon rank sum test implemented by the *FindAllMarkers()* methods of the Seurat package. This was performed before and after correcting for the transcriptome alterations induced by enzymatic digestion on dura meningeal ILCs. To do this correction, we generated a list of genes reported to be associated to enzymatic digestion (Sup. Table 3)^32,68,70^ and removed them from the differentially expressed genes found in dura meningeal ILCs before subsequent analysis. Differential gene expression analysis between compartments for the whole dataset or per cluster was done after randomized subsetting and repeated several times to confirm consistency in the results. Furthermore, compartment-specific differential analysis was done at the single cell level and at pseudobulk level, integrating both analysis as a final step for the selection of the compartment-specific differentially expressed genes. At the single cell level, the *FindAllMarkers()* function from the Seurat package based on Wilconxon test (logfc.threshold= 0.2, only.pos =FALSE, min.pct = 0.1) was used, while the Limma package functions *lmFit(), eBayes(), topTable()* were used to implement empirical Bayes linear models for identifying DEGs on the transformed pseudobulk data. Pseudobulk data was generated by converting the Seurat object into a SingleCellExperiment object, and applying the *aggregateAcrossCells()* function. The Fisher’s method for combining p-values was then used on the DEGs with a fold-change above or below 0.8/-0.8 in both analysis and DEGs were selected based on Fisher p value < 0.05.

### Pathway enrichment

Overrepresentation enrichment analysis with the function enricher from the *clusterProfiler* package (v4.6.2; Wu et al.,(2021))^71^ was used on the cluster-specific and compartment-specific differentially expressed genes to determine significantly enriched Hallmark (p<0.05) and Gene Ontology (GO) terms (adj.p < 0.05 and k.K (significant genes in set/total genes in set) > 0.1). In addition, the fgsea package (v1.24.0; Korotkevich et al., (2016))^72^ for fast pre-ranked gene set enrichment analysis (GSEA) was used on the pre-ranked gene lists generated, based on the fold-change values calculated by the *FindMarkers* function using the Wilcoxon rank sum test.

### Trajectory analysis

Monocle 3 (v1.3.4)^73,74^ was used to analyze plasticity related trajectories between ILC3 and ILC1 clusters. The function *as.cell_data_set()* was used to convert the Seurat object into a Monocle3 object. After graph learning was performed (*learn_graph()*), we ordered the cells with the function *order_cells(),* setting the starting node embedded in the NKp46^+^ ILC3 cluster. Trajectories were then visualized using the *plot_cells()* function. We then identified genes co-regulated along the pseudotime with the graph-autocorrelation function graph_test(neighbor_graph= “principal_graph”) and plotted the top 8 most significant ones in graphs where cells were ordered onto a pseudotime trajectory based on these ILC3 and ILC1- identity markers.

### Analysis of CNS-derived CD45^+^ immune cell scRNA-seq data

Data from whole brain and micro-dissected CNS barrier-sorted immune cells generated by Van Hove, et al. (2019)^32^ was downloaded and analyzed using Seurat standard workflow (see above). ILC clusters were identified by excluding clusters with expression for other CNS-immune cells and selecting those with the highest ILC gene expression score (depicted in Sup. Table 1).

### Statistical analysis

GraphPad Prism 9 was used for statistical analysis. Whitney test for two group comparisons and Kruskal-Wallis test with Dunn’s for multiple comparison were used for statistical analysis. P values of p > 0.05 were considered as not significant, P values were considered as followed: *p < 0.05, **p < 0.01, ***p < 0.001. Figures show bars or dots indicating mean ± SEM unless stated. Due to the exploratory nature of the study, a power calculation was not performed to determine the sample size of each group. Therefore, a rule of minimum three animals to maximum eight animals per group was set for data sampling. Data analysis was not performed blindly to the experimental conditions.

### Data analysis and visualization

Data was analysed using several specialized tools depending on the technique (Flowjo, Graphpad Prism, R v5.0.2, ImageJ) and figures were created and edited in Adobe Illustrator.

## Data availability

The authors declare that the data supporting these findings are present in the paper or the Supplementary Materials. Raw data are available from the authors upon reasonable request. RNA sequencing data will be available through GEO #[submission in progress].

## Author contribution

A.D.R.S. designed the study, performed the experiments with the help of B.L.H. and E.V., analyzed the data, prepared the figures, and wrote the manuscript. C.S., O.H., A.D., and C.R. provided technical and conceptual advice and relevant resources. CI-D. designed and directed the study and wrote the manuscript. All authors fully qualify for authorship and have approved the final version of the manuscript.

## Ethics declaration

The animal study was reviewed and approved by Berlin State Office for Health and Social Affairs (LAGeSo), Berlin, Germany.

## Conflict of interest

The authors declare that the research was conducted in the absence of any commercial or financial relationships that could be construed as a potential conflict of interest.

### Acknowledgement

We thank all the members of the Infante-Duarte lab for their helpful experimental advice and discussions. We especially thank Natascha Asselborn for all her technical assistance and Thordis Hohnstein with her support establishing the new transgenic animal lines in our breeding barrier. We also thank Single Cell Discoveries for their help with single cell sequencing services. In addition, we would like to thank Franke Vedran for his expert advice and guidance during the analysis of the single cell sequencing data. We would like to also acknowledge Caroline Braeuning from the BIH/MDC Genomics Technology Platform for her expert technical assistance and services in our FACS sorting experiments. We thank the Advanced Medical BioImaging Core Facility of the Charité-Universitätsmedizin Berlin (AMBIO) for support in acquisition of our imaging data and the Research Institute for Experimental Medicine (FEM) of the Charité (Berlin, Germany) for their excellent maintenance of our animal colonies and for the careful control of our breeding to provide us with animals along all the developmental timepoints studied in this paper. This study was funded by the German Research Foundation (Deutsche Forschungsgemeinschaft, DFG), Projekt Nr. 534335032, Grant ID: IN 156/8-1, and by the Hertie Foundation (GHS, Gemeinnützige Hertie-Stiftung), through the Hertie Network of Excellence in Clinical Neuroscience (Project Nr. P1200003). CID was also supported by the DFG-funded SFB1340-2, Grant Nr. 372486779, Project B05.

