## Supplementary figures and figure legends for "Perinatal brain group 3 innate lymphoid cells are involved in the formation of murine dural lymphatics"

#### Supplementary figure legends

**Supplementary Figure 1. scRNA-seq of young adult CNS compartments show similar profile of resting CNS tissue resident ILCs. (A)** Complete flow cytometry gating strategy for identification of ILC in adult CNS WT young adult (8-11 weeks old) mice. **(B)** UMAP visualization of the scRNA-seq from total CD45$^+$ immune cells found within whole brain and micro-dissected CNS barriers (dura mater and choroid plexus). Data are from Van Hove et al., (2019). **(C)** Violin plot represent the distribution of ILC gene expression programs, defined in table 1, grouped by clusters. **(D)** UMAP representation of the lymphocyte subclusters (left), and feature plot of T cell associated markers expressed among the new obtained clusters (right). **(E)** Heatmap displaying the top 15 DEGs among ILC-specific clusters (clusters 1, 2 and 5). Color scale represents average expression. **(F)** Violin plots of lineage-specific cytokines and Ki67. **(G)** Representative gating for identification i.v. CD45-PE$^+$ immune cells (CD45-PerCP$^+$) in peripheral and CNS compartments. Quantification is represented as mean ± SD, n= 6. **(H)** UMAP plot representation showing average expression of circulating and tissue resident associated markers. **(I)** Dotplot of selected gene expression within clusters. Color scale represents average expression, dot size visualizes fraction of cells within the cluster expressing the gene. **(J)** On the left is the representative gating strategy for the identification of ST2$^+$ ex-ILC3s using the Rorc(gt)$^{cre/wt}$×R26$^{eYFP}$ FM$^+$ mice 2–3-month-old. On the right, the graph depicts the proportion of RORgt$^-$YFP$^+$ ex-ILC3s among ST2$^+$ ILCs in brain and dura mater. Graph is depicted as mean ± SEM., n = 9 examined over 2 independent experiments.

**Supplementary Figure 2. No gender differences in the CNS ILC compartment of young adult mice. (A)** Quantification of the absolute numbers of each ILC subset and the percentage over total ILCs in whole brain and **(B)** dura mater of male and female young adults (8-10 weeks old). Graphs are depicted as mean ± SEM., n = 6-8 / group. Dura mater were pooled in groups of two when necessary. **(C)** Quantification of Ki67 expression among ILCs within the different CNS compartments in male and female mice. Data are depicted as ± SEM., n = 3 / group. Data were analyzed by a Mann Whitney t-test.

**Supplementary Figure 3. Eomes$^-$ and Eomes$^+$ cells are present in the brain along ontogeny. (A)** Quantification and representative plots of NKp46 and Eomes expression by CD122$^+$T-bet$^+$ cells along ontogeny. **(B)** Frequency and **(C)** absolute numbers of group 1 ILC subsets identified as Eomes$^-$ (ILC1) and Eomes$^+$ (NK) in the brain along ontogeny. Data are depicted as mean ± SEM., n = 3-7 / timepoint examined over ≥ 2-3 independent experiments. **(D)** Quantification of Ki67 expression in Eomes$^+$ and Eomes$^-$ in brain along ontogeny.

Quantification as mean ± SEM., n = 4-8 / timepoint. Data are representative of ≥ 3 independent experiments. From E16 to P1 whole litter were pooled. From P9 to P15 onwards, animals were pooled in 2-3 when necessary.

**Supplementary Figure 4. scRNA-seq analysis of P9.5 brain and dura ILCs identified presence of all ILC subsets. (A)** Heatmap displaying the top 15 differentially expressed genes (DEGs) within the identified ILC clusters comparing brain versus dura cells. **(B)** Dot plot and **(C)** UMAP plot representation showing average expression of other brain immune cell, common lymphoid progenitor (CLP) and ILC progenitor (ILCp) cell markers, respectively. **(D)** Violin plot visualization of the expression of effector cytokines within the identified helper-ILC clusters in brain and dura meninges.

**Supplementary Figure 5. Transitory presence of ILC3s during brain ontogeny. (A)** Absolute numbers of $CD4^{+/-}CCR6^{+}$, $NKp46^{+}$ and double-negative (DN) $ROR\gamma t^{+}$ ILC3 subsets (n = 3-8 / timepoint, mean ± SEM.). Data are representative of ≥ 3 independent experiments. **(B)** Heatmap displaying the top 15 DEGs as assessed by scRNA-seq within ILC1/ILC3 subclusters. **(C)** Violin plot of selected markers expression within ILC1/ILC3 subclusters. **(D)** Percentage from total analyzed cells of each identified subcluster among ILC1 and ILC3s in whole brain and dura. **(E)** Quantification of Ki67 expression in each ILC3 subset in brain along ontogeny (n = 3-7 / timepoint). From E16 to P1. whole litters were pooled, from P9 to P15 onwards. 2-3 animals were pooled if needed. Data are depicted as mean ± SEM.

### Extended data

#### Supplementary Figure 1

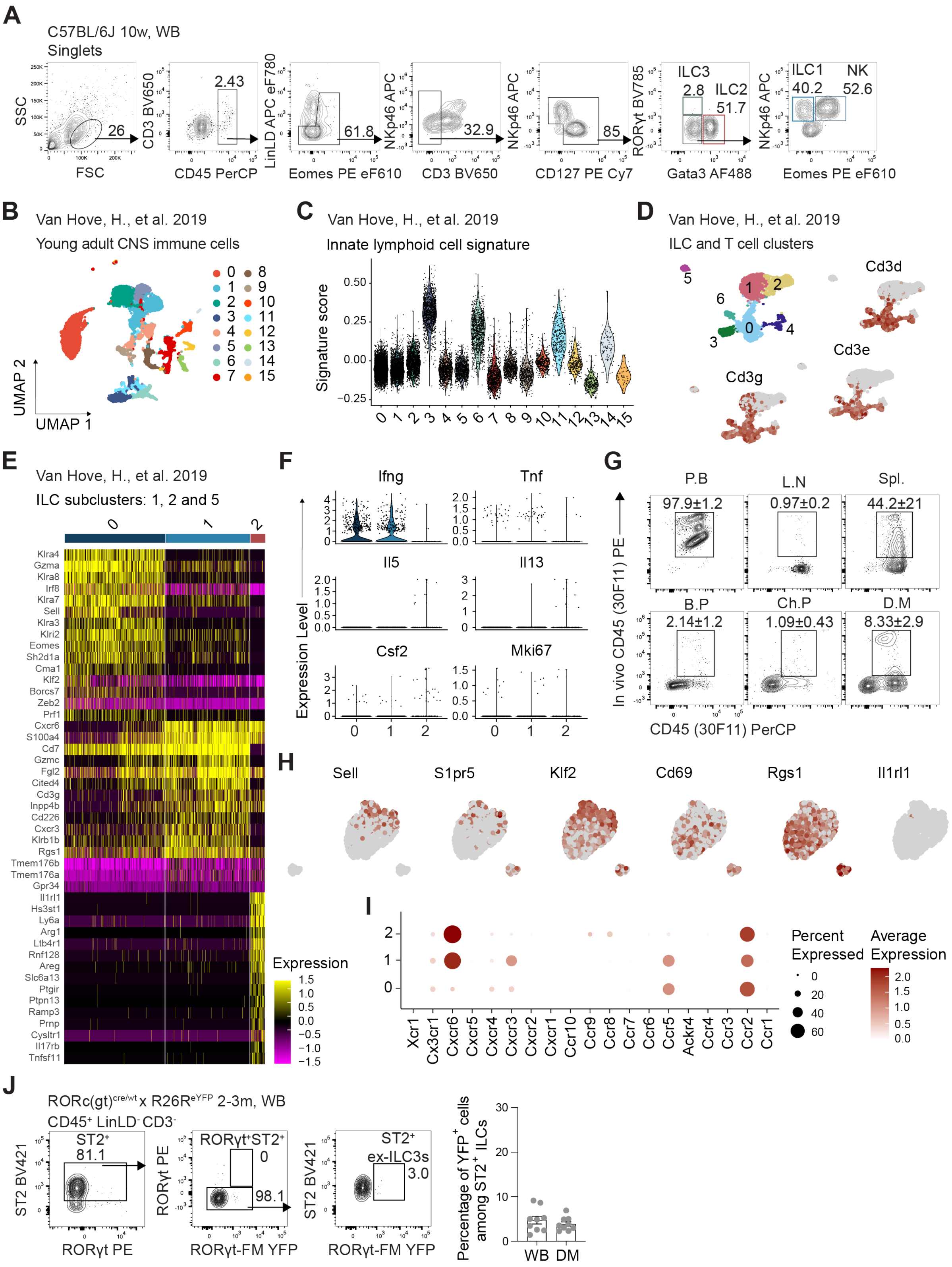

Supplementary Figure 2

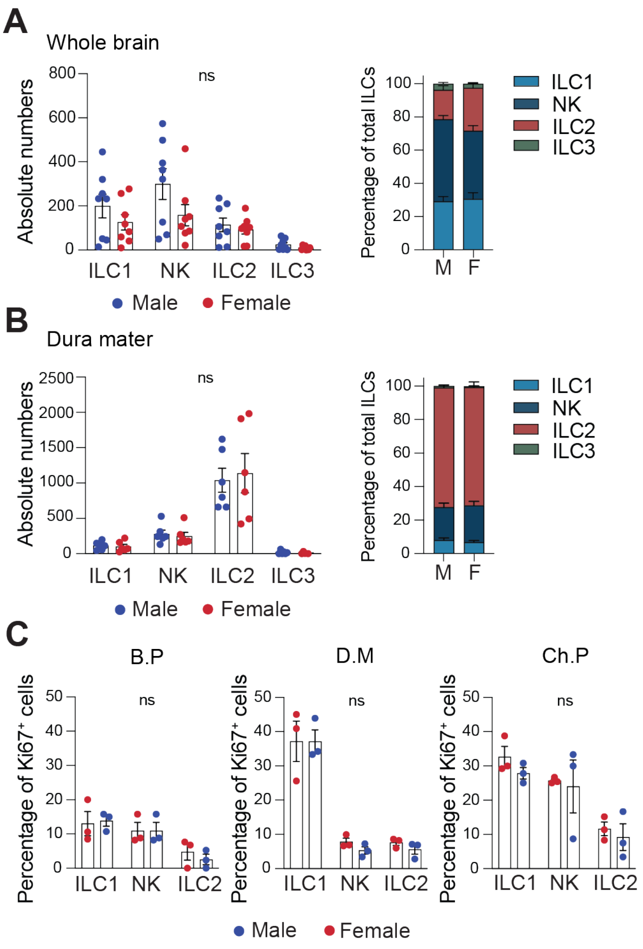

Supplementary Figure 3

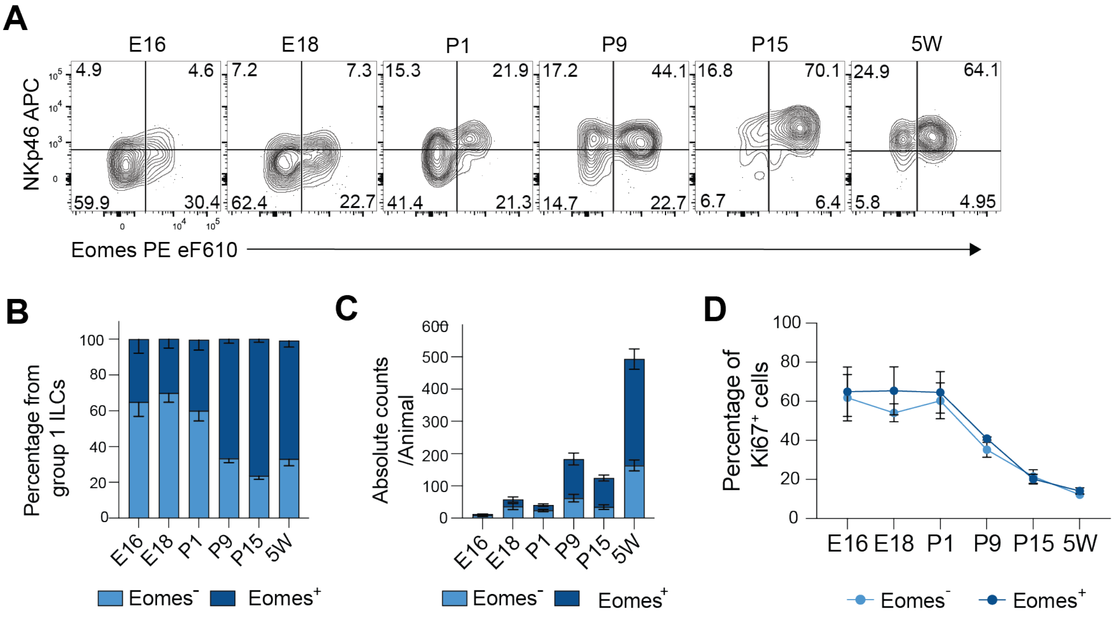

Supplementary Figure 4

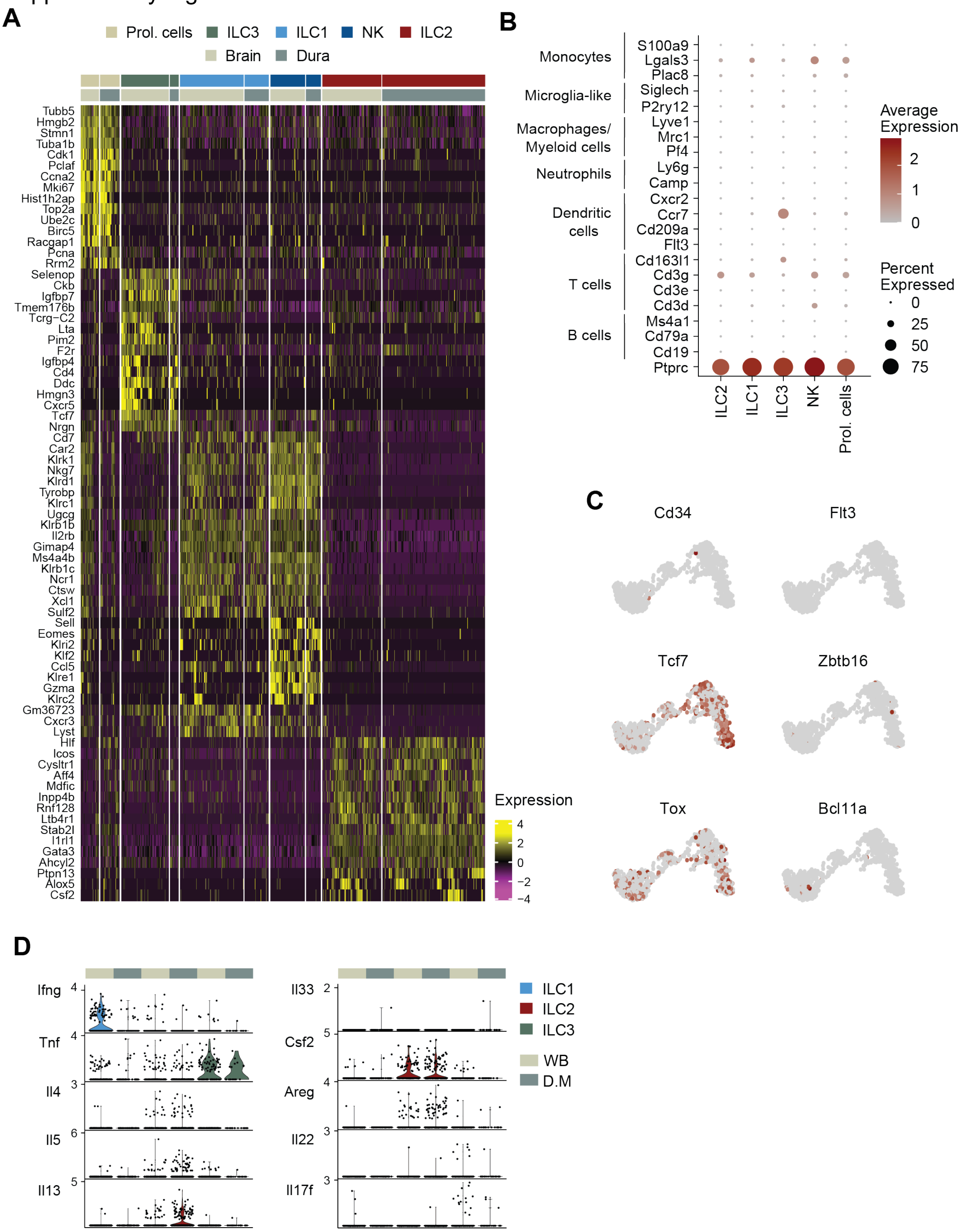

Supplementary Figure 5

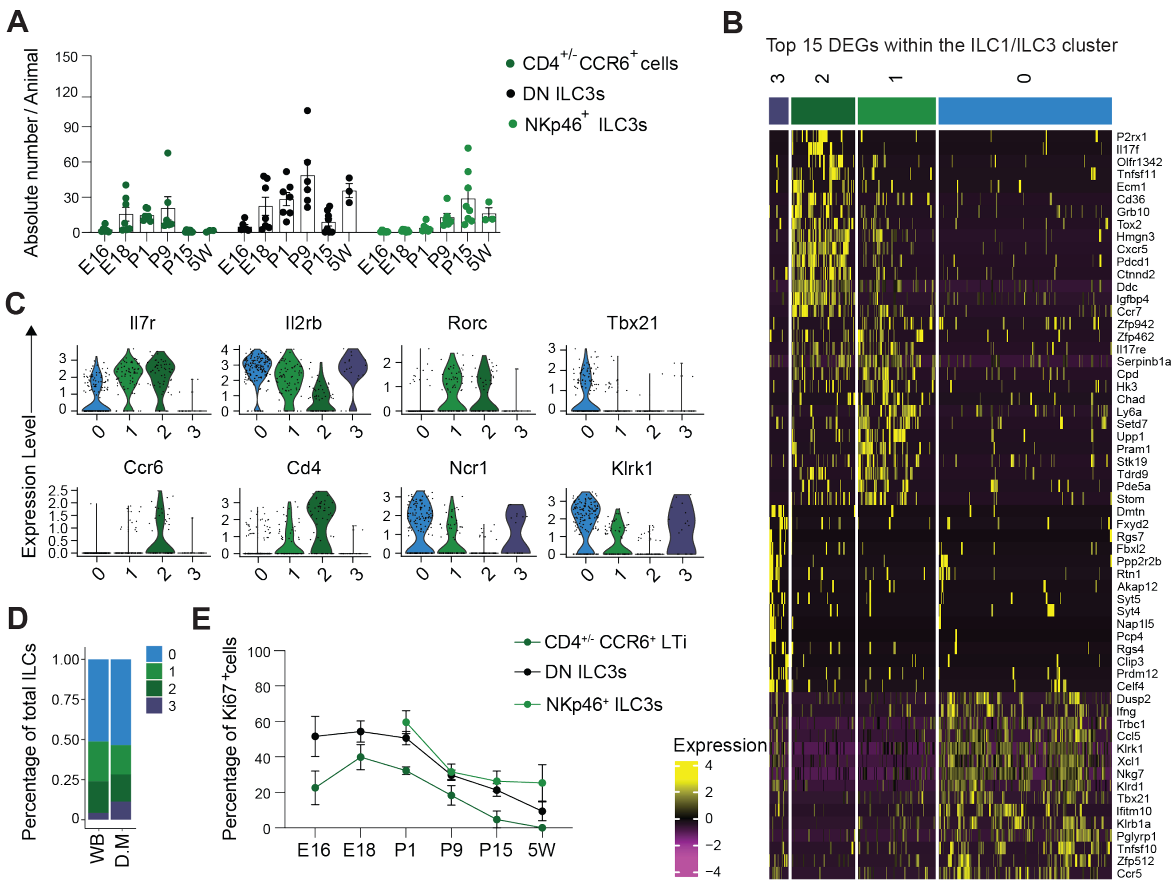
